# OmniRE-seq: A Dual-Method Framework for Decoding RNA Editing Dynamics in Disease and Therapy

**DOI:** 10.64898/2026.10.22.760919

**Authors:** Yuqing Hang, Hao Liu, Wangchang Li, Pinbo Liu, Jihong Yang, Yuxiang Zhao

## Abstract

RNA editing is a widespread post-transcriptional modification involved in transcriptome diversity and disease regulation. However, its quantification remains technically challenging. We present OmniRE-seq, a dual-framework combining RE-seq for bulk and scRE-seq for single-cell RNA-seq data, enabling multi-level quantification, differential analysis, and functional annotation of RNA editing. Using RE-seq, we profile editing alterations across diverse contexts -- including murine sepsis, MDS, drug-treated melanoma, and colorectal cancer -- demonstrating the utility of locus-, gene-, and burden-level metrics in capturing biologically meaningful differences. scRE-seq further resolves cell-type-specific editing patterns by aggregating cells of shared identity into pseudo-bulk units (Spots), enhancing editing signal while preserving cellular resolution in colorectal cancer. Integrated bulk and single-cell analysis identified convergent editing signals and therapeutic targets, exemplified in a mouse model of thymic atrophy treated with *Ganoderma lucidum* spores. Together, OmniRE-seq offers a standardized and scalable toolkit for dissecting RNA editing landscapes across resolution levels, advancing both mechanistic discovery and translational application in precision medicine.

## Introduction

RNA editing, a ubiquitous post-transcriptional mechanism in eukaryotes, modifies mRNA nucleotide sequences through transcriptional inaccuracies or post-transcriptional modifications ^1–4^. First documented in 1986, this process remains underexplored in translational contexts despite its biological significance ^5^. Catalyzed primarily by deaminases, RNA editing dynamically regulates gene expression and protein function by converting specific nucleotides ^6^. The most prevalent forms involve cytosine-to-uracil (C-to-U) editing by the APOBEC enzymes and adenosine-to-inosine (A-to-I) editing by the ADAR proteins, collectively representing ~90% of known editing events ^7–9^. Rarer forms (e.g., C-to-G, C-to-A), lack identified mediators, underscoring gaps in mechanistic understanding ^10^.

Unlike heritable DNA mutations, RNA editing enables transient, adaptive responses to environmental and pathological stimuli ^11^. The primary biological function of RNA editing is its ability to alter amino acid sequences, thereby affecting protein coding ^12^. Beyond protein recoding, RNA editing also involves modifications in non-coding regions, which can alter RNA secondary structures, induce RNA splicing events, and target gene expression regulation through edits in the 3′ UTR and miRNAs ^13^. These multifaceted roles position RNA editing as a critical player in disease pathophysiology ^14,15^.

Emerging evidence implicates dysregulated RNA editing in onset and progression of diverse human diseases, underscoring its dual relevance as a diagnostic biomarker and therapeutic modulator ^16–18^. In cancer, aberrant RNA editing perturbs genes related to apoptosis, metastasis, and therapy resistance, while mutations in APOBEC and ADAR enzymes alter oncogene and tumor suppressor expression, promoting malignant survival under stressful conditions ^19^. In neurodegenerative diseases, such as Alzheimer’s disease and amyotrophic lateral sclerosis (ALS), impaired A-to-I RNA editing disrupts glutamatergic signaling and synaptic integrity, contributing to neuronal vulnerability and disease progression ^20^. Moreover, in autoimmune and inflammatory disorders, RNA editing has been shown to modulate immune receptor expression and cytokine signaling pathways, influencing both innate and adaptive immune responses ^21^. Notably, while most therapeutic agents are not designed to directly modify RNA sequences, pharmacological or environmental perturbations can influence RNA editing patterns indirectly—via changes in editing enzyme activity, RNA substrate structure, or cellular signaling—resulting in dynamic, reversible transcriptomic alterations. Capturing such adaptive editing responses is crucial for understanding both physiological regulation and therapeutic impact. Collectively, these findings establish RNA editing as both a disease-relevant modulator and a potential source of diagnostic and therapeutic targets ^22^.

Advances in high-throughput sequencing have enabled genome-wide editing analyses, yet technical hurdles persist. Distinguishing true RNA editing events from sequencing artifacts, mapping errors, and genomic variants (e.g., SNPs) remains challenging ^23–25^. While integrating whole genome or exome sequencing and SNP databases mitigates false positives, cost and accessibility constraints limit utility. Current bioinformatics tools further exhibit biases toward A-to-I events, neglecting rare editing types ^26,27^. Quantification challenges compound these issues. Current pipelines lack standardized methods for cross condition comparisons, often overlooking biological variability and sample heterogeneity ^28^. Bulk RNA-seq obscures cell-type-specific editing patterns, whereas single-cell approaches grapple with low sequencing depth and technical noise ^29^. This gap impedes exploration of RNA editing’s spatiotemporal dynamics and its role in cellular heterogeneity—a critical frontier for understanding disease progression.

To address these limitations, we introduce OmniRE-seq, a comprehensive framework for detecting, quantifying, and functionally analyzing RNA editing events in both bulk RNA-seq and scRNA-seq data (**Figure 1**). This unified pipeline synergizes preprocessing, variant calling, annotation, and functional characterization, thereby resolving RNA editing events with high precision. By integrating population-wide and single-cell-level insights, OmniRE-seq advances the exploration of RNA editing’s role in disease pathogenesis and precision medicine. Our framework overcomes prevailing technical challenges, enabling robust, context-aware quantification of editing dynamics. This capability not only enhances mechanistic studies but also accelerates the translation of RNA editing discoveries into clinical biomarkers and targeted therapies.

**Figure 1.**
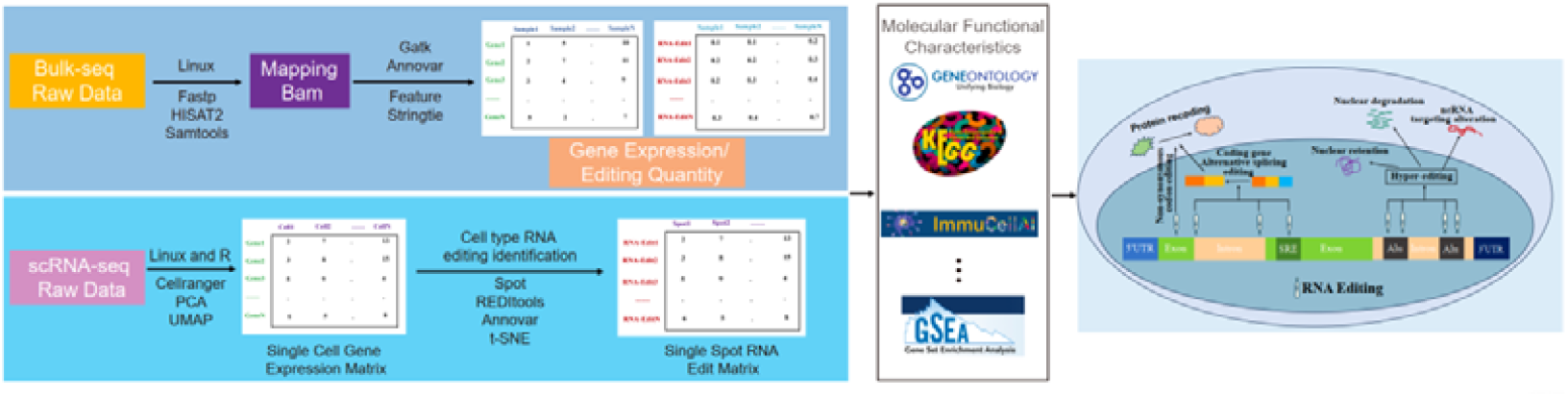
Workflow diagram of an integrated dual-method framework for bulk and single-cell RNA-Seq-based RNA editing quantification.

## Results

### RE-seq framework for bulk RNA editing analysis

The absence of standardized RNA editing quantification frameworks has hindered robust cross-study comparisons. To address this gap, we present the RE-seq pipeline, a computational framework for systematic identification, quantification and differential analysis of RNA editing events in bulk RNA-seq data (**Figure 2**). The pipeline integrates upstream RNA-seq data processing, editing site detection, quantitative characterization at multiple biological scales, differential editing analysis, and downstream function interpretation to capture the potential biological relevance.

**Figure 2.**
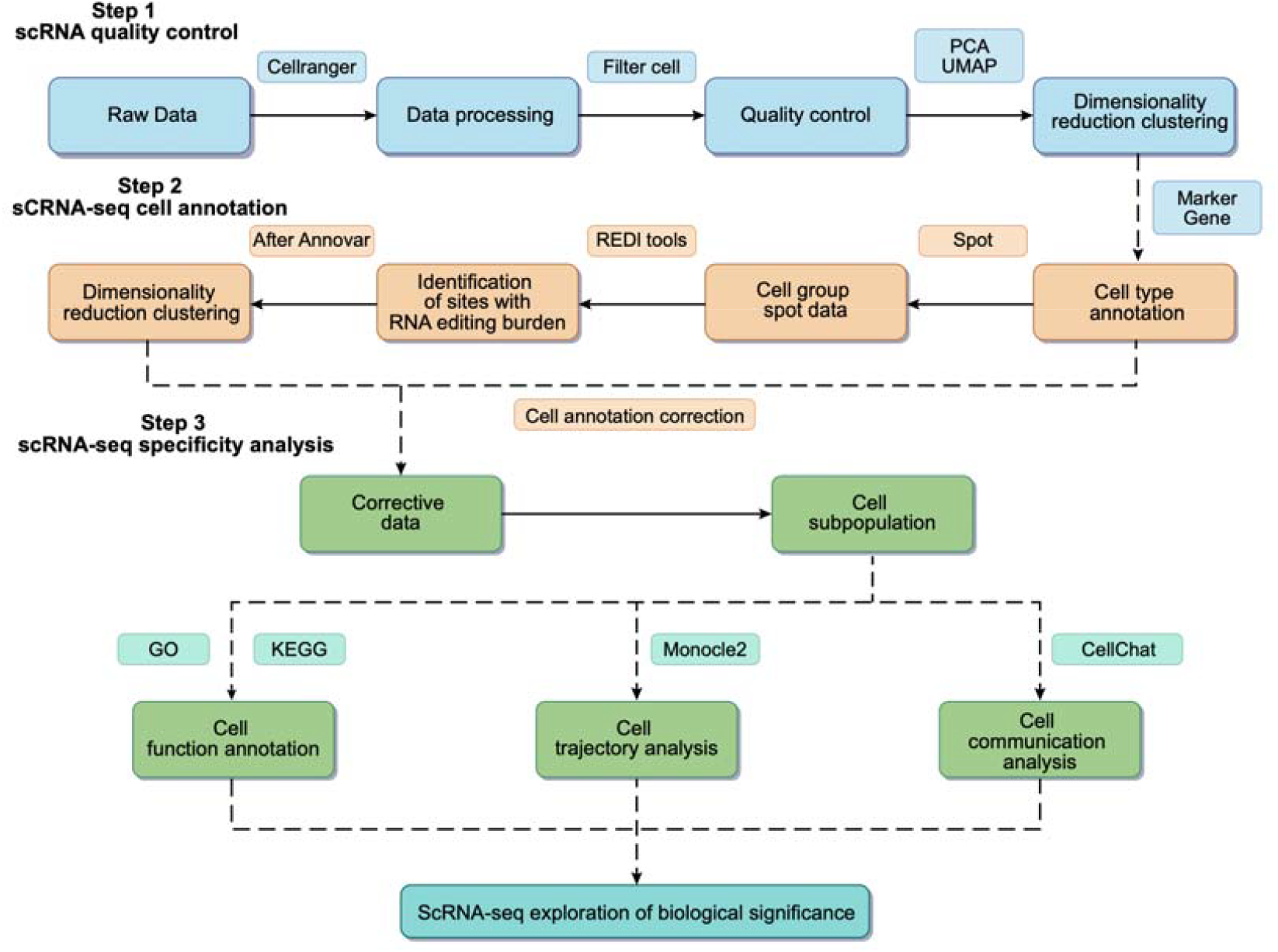
Workflow diagram illustrating an integrated framework for RNA editing quantification and analysis in bulk RNA-Seq data.

In the first step, raw sequencing reads are preprocessed using fastp ^30^ for quality control, followed by alignment to the reference genome using HISAT2 ^31^ and format conversion and BAM processing with SAMtools ^32^. High-quality BAM files are used for transcript reconstruction (StringTie) ^33^ and variant calling (GATK) ^34^, enabling identification of candidate RNA editing-like sites. Identified variants are then annotated using ANNOVAR ^35^, providing functional context for each editing event.

A defining strength of RE-seq is its multi-tiered quantification system, which evaluates RNA editing activity across three biological scales. At the **loci level**, editing is further quantified using Loci Editing Read Counts (LERC), Loci Editing Frequency, and RNA Editing Loci Burden (RELB), enabling the precise characterization of site-specific editing dynamics. At the **gene level**, Gene Editing Read Counts (GERC), Gene Editing Frequency, and RNA Editing Gene Burden (REGB) aggregate editing activity across entire genes, facilitating gene-centric analyses. At the **sample-level** metric, the RNA Editing Burden (REB) number represents the total count of high-confidence RNA editing-like sites detected per sample, while REB normalizes the REB number by the total length of annotated protein-coding regions, offering a global measure of editing activity that supports cross-condition comparisons. To evaluate differential editing between biological conditions, RE-seq adopts data distribution-specific strategies. Count-based metrics (LERC, GERC) are analyzed using DESeq2 ^36^ and limma ^37^, while burden-based metrics (RELB, REGB), which deviate from standard parametric assumptions, are tested using a permutation-based framework ^38^. Beyond differential quantification, RE-seq incorporates a suite of downstream analyses to interpret the biological implications of RNA editing. Functional enrichment analysis is performed using Gene Ontology (GO) and Kyoto Encyclopedia of Genes and Genomes (KEGG) pathways, applied to both differentially edited genes and differentially expressed transcripts. To evaluate the immunological relevance of RNA editing, RE-seq integrates immune cell deconvolution approaches, quantifying immune infiltration levels across samples and associating them with editing burden and gene-level changes ^39^.

To further elucidate the regulatory impact of RNA editing, RE-seq incorporates a multi-omics integration module. This framework supports joint analysis between RNA editing and other molecular layers, such as gene expression and genotypic variation. Specifically, it includes quadrant-based classification of editing–expression relationships and the identification of editing quantitative trait loci (edQTLs) when matched genotype data are available. By unifying editing, transcriptomic, and immune features, RE-seq enables the discovery of biologically meaningful editing events and their potential downstream regulatory associations within disease-specific regulatory networks. This integrative approach provides a robust and scalable framework for decoding the biological significance of RNA editing in bulk RNA-seq data, facilitating the discovery of RNA-editing-associated pathways and candidate biomarkers of therapeutic-associated editing events for future validation.

### scRE-seq pipeline for single-cell RNA editing analysis

While bulk RE-seq has advanced our understanding of tissue- and disease-level RNA editing variation, resolving cell-type-specific editing dynamics remains critical for dissecting disease heterogeneity. To address this limitation, we develop scRE-seq, a pipeline optimized for analyzing RNA editing patterns in 10x Genomics single-cell RNA-seq (scRNA-seq) data. As depicted in **Figure 3**, the scRE-seq pipeline integrates three core modules: (1) preprocessing and quality control, (2) cell-type annotation, and (3) cell-type-specific differential RNA editing analysis.

**Figure 3.**
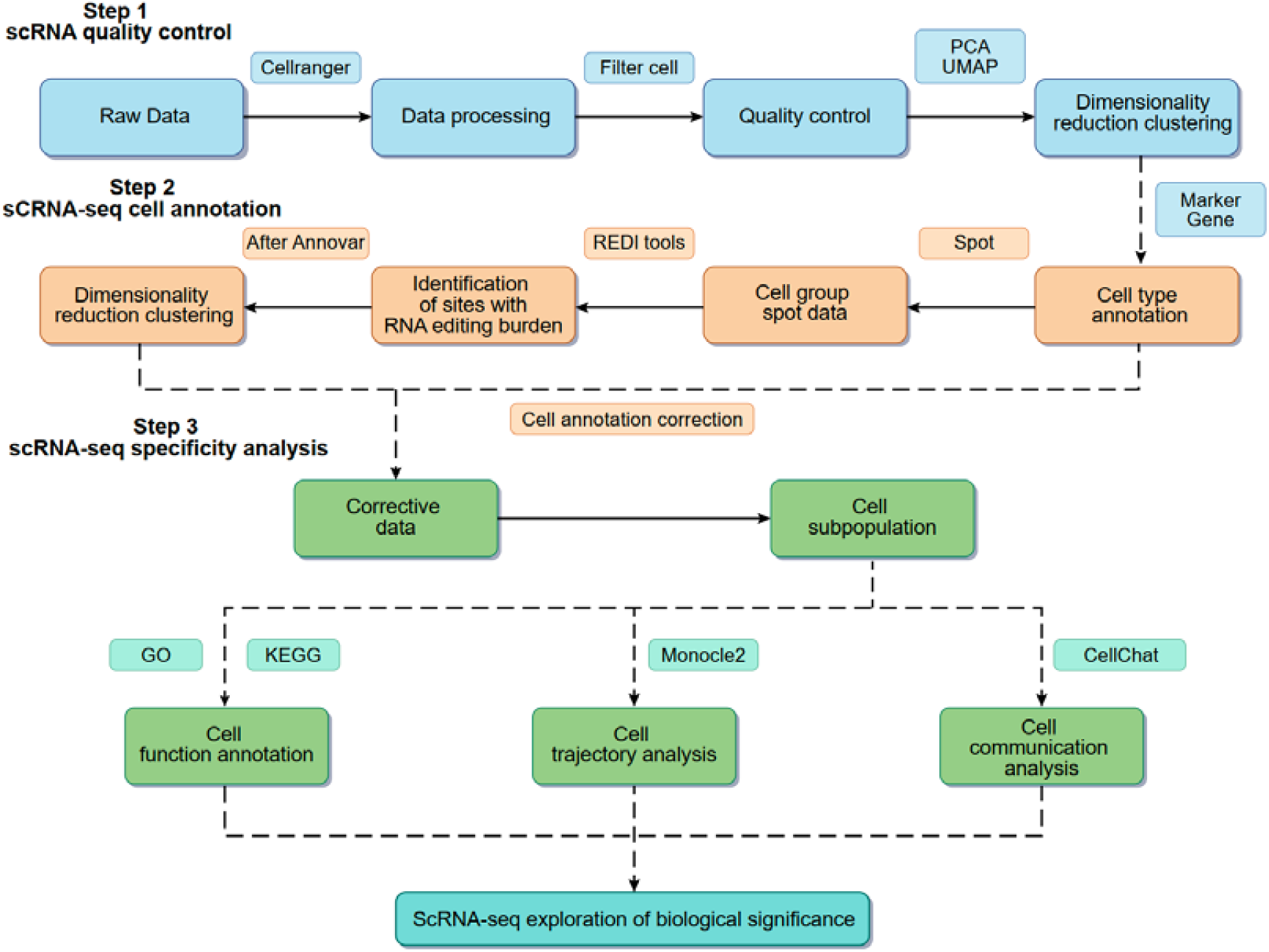
Workflow diagram illustrating an integrated framework for RNA editing quantification and analysis in single-cell RNA-Seq data.

The data processing and dimensionality reduction steps in scRE-seq align with conventional scRNA-seq pipeline. However, single-cell RNA editing analysis faces challenges due to sparse transcriptome coverage and limited sequencing depth inherent to 10x Genomics data. To mitigate these limitations, scRE-seq incorporates multiple optimizations. First, we introduce Spots, pseudo-bulk units constructed by aggregating cells from the same donor and annotated cell type to increase effective read deapth for editing detection (**Figure 4A**). Second, we implement a refined PCR duplicate removal strategy, defining duplicates strictly as reads sharing identical cell barcodes (CBs) and unique molecular identifiers (UMIs), ensuring accurate estimation of editing events (**Figure 4B**). Benchmarking across Spot sizes ranging from 10 to 100 cells revealed that 20-cell pools optimally balanced editing detection sensitivity measured by event yield, annotation depth, and cellular resolution evaluated via Davies–Bouldin Index (DBI), K-nearest neighbors score (KNN), and Silhouette coefficient (**Figure 4C, Figure S1**). This configuration enables robust quantification of RNA editing dynamics while preserving intra-sample cellular architecture.

**Figure 4.**
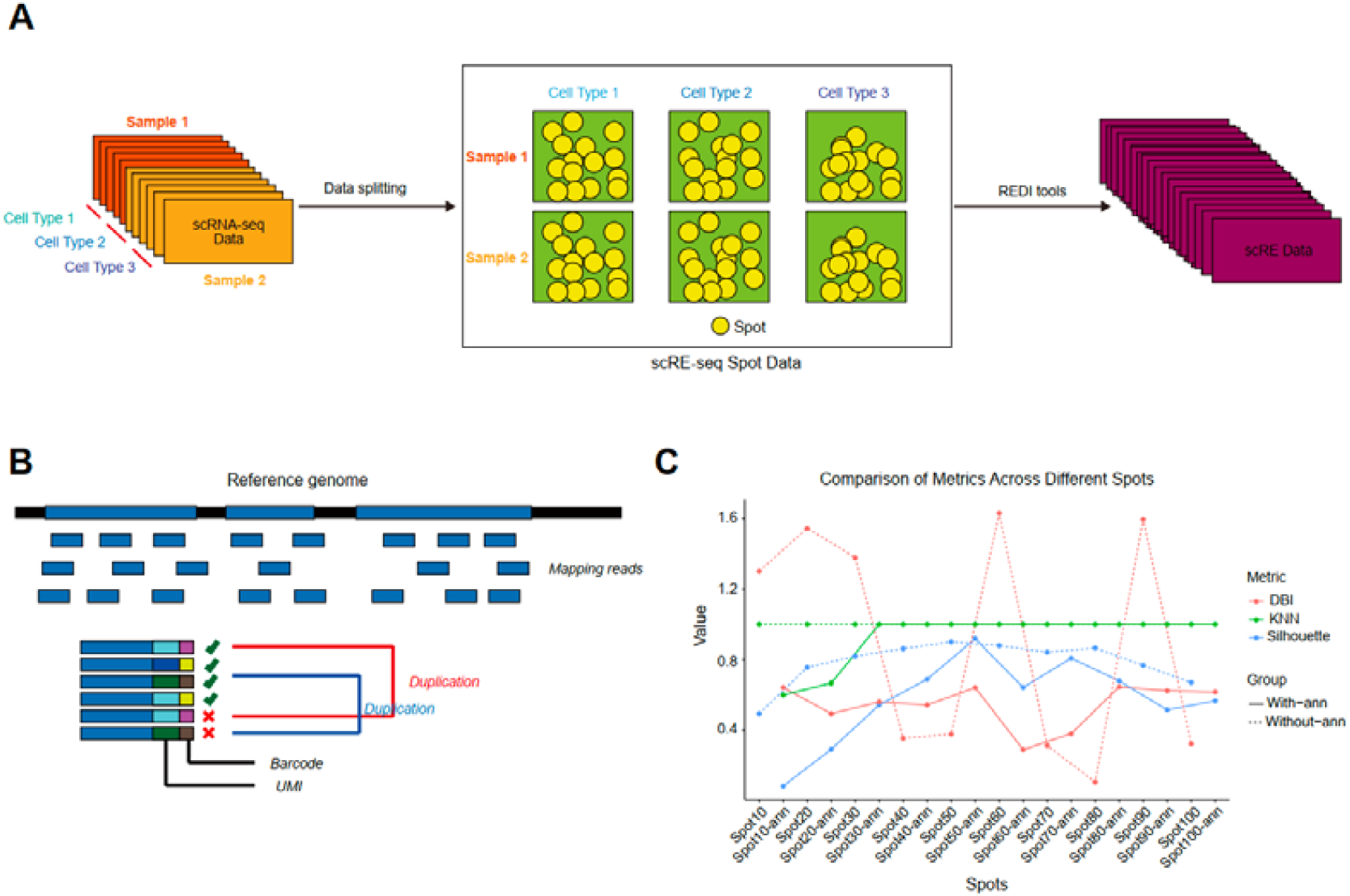
Workflow diagram for spot-based pooling optimization in single-cell RNA editing detection. (A) Schematic of spot construction. (B) Strategy for PCR duplicate removal. (C) Optimal threshold filtering of spot sizes.

To improve cell-type annotation accuracy, we implement an RNA-editing-guided quality control strategy. Using scRNA-seq derived cell-type labels as reference, we cluster Spots based on RNA editing profiles and evaluate cluster purity. Clusters with >40% of Spots from divergent cell types are flagged as low-confidence and exculed from downstream differential editing analysis. This filtering reduces potential ambiguous or mixed clusters, particularly low-abundance cell populations such as endothelial, NK, mast cells, and neutrophils. Multimodal clustering post-QC shows improved concordance with known cell types, validating the refinement strategy (**Figure S2**).

High-confidence editing sites are then subjected to downstream analyses. Functional enrichment, pseudotime trajectory inference, and intercellular communication modeling reveal the regulatory potential of RNA editing across differentiation states and cell–cell interaction networks. These analyses offer a comprehensive map of RNA editing heterogeneity and its functional implications at single-cell resolution.

### RNA editing in Sepsis-induced myocardial dysfunction in a mouse model

To explore the role of RNA editing in sepsis-related cardiac injury, we applied the RE-seq framework to myocardial tissue from a murine sepsis model, comprising control, sepsis, and sepsis-treatment groups. Editing events were quantified using the gene-level REGB metric to characterize gene-level edtiting alterations.

The global distribution of RNA editing types was dominated by canonical C-to-U and A-to-I conversions, which together accounted for approximately 70% of all events (**Figure 5A**). These edits predominantly occurred in intronic and exonic regions and were mostly unannotated, though a substantial proportion included nonsynonymous substitutions, indicating potential coding impact (**Figure S3A-B**). Principal component analysis (PCA) based on REGB clearly distinguished these three groups, underscoring transcriptome-wide editing shifts associated with sepsis and therapeutic intervention (**Figure 5B**). Consistent with this, total REB was significantly elevated in septic and treatment groups relative to controls (**Figure S3C**), supporting a sepsis-induced increase in editing activity.

**Figure 5.**
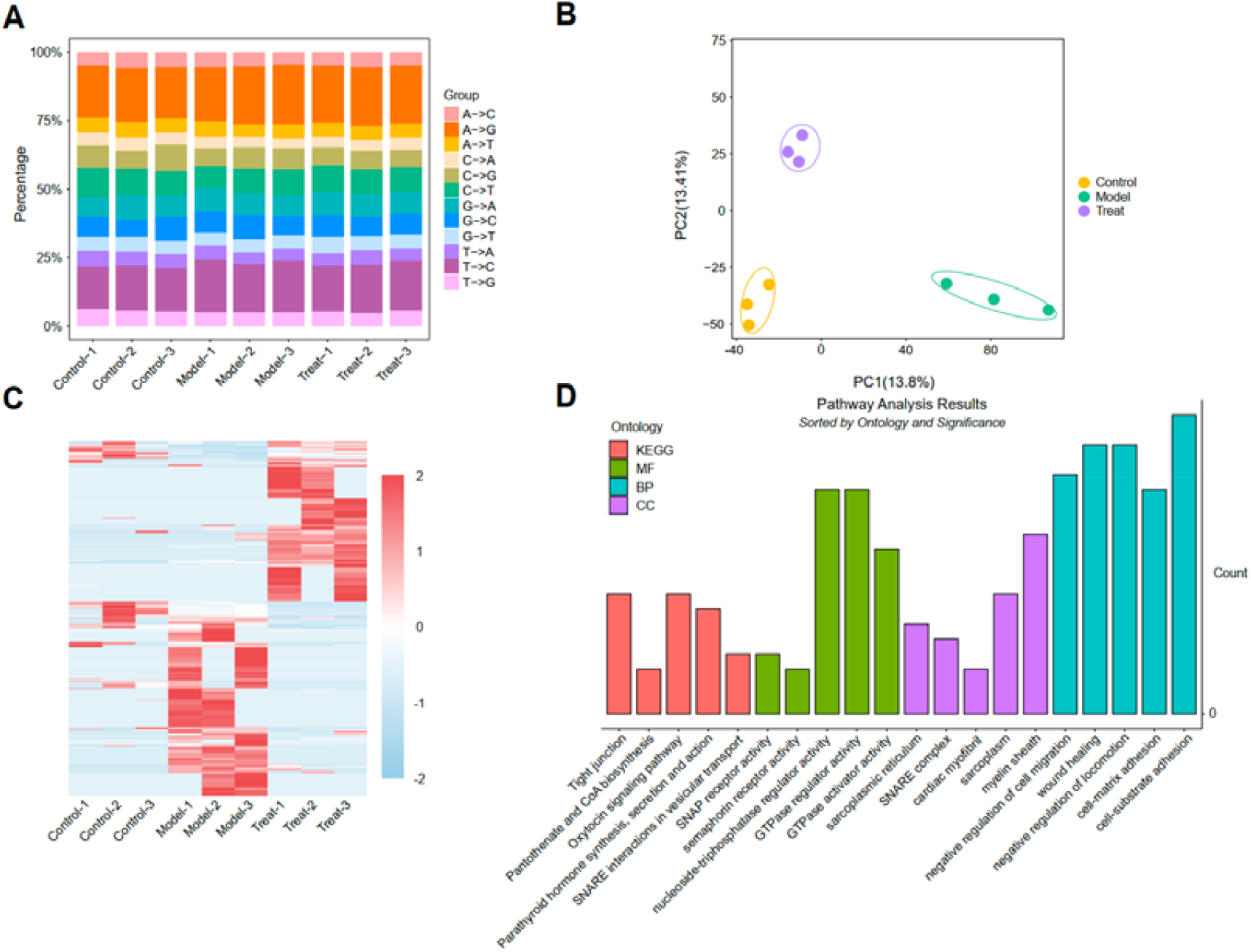
The landscape of RNA editing in sepsis-induced myocardial dysfunction in a mouse model. (A) Proportion of RNA editing types across the three experimental groups (Control, Sepsis, and Sepsis Treatment). (B) PCA of RNA editing-like events based on REGB metric. (C) Heatmap of differentially edited genes using REGB metric. (D) GO enrichment of differentially edited genes.

Differential editing analysis based on REGB identified a set of significantly altered genes, whose editing profiles clustered distinctly across samples (**Figure 5C, Figure S3D**). Functional enrichment of these genes revealed strong associations with immune signaling and inflammatory regulation. GO analysis implicated pathways involved in T-cell receptor signaling, innate immune response, and inflammatory processes. KEGG pathway analysis further highlighted T-cell receptor signaling, hematopoietic cell lineage, and viral myocarditis, while Gene Set Enrichment Analysis (GSEA) identified IL-17 signaling as a potentially druggable axis of regulation in sepsis (**Figure 5D, Figure S3E**). Collectively, these results demonstrate widespread RNA editing alterations during sepsis progression and therapeutic response, highlighting RE-seq’s utility in uncovering editing-associated regulatory programs in inflammatory disease models.

### RNA editing in the Myelodysplastic Syndromes (MDS)

To investigate the role of RNA editing in the pathogenesis of Myelodysplastic Syndromes (MDS), we applied the RE-seq framework to bone marrow RNA-seq data from patients categorized into high-risk (MDS-H), low-risk (MDS-L), and control (MDS-C) groups. RNA editing activity was quantified using the GERC metric to capture gene-level alterations.

Global profiling revealed that the editing landscape was dominated by canonical A-to-I and C-to-U substitutions, with the majority of events localized in intronic and exonic regions (**Figure 6A, Figure S4A**). Functionally, annotated editing events were enriched for synonymous and nonsynonymous substitutions, with potential impacts on protein coding (**Figure S4B**). PCA of GERC values revealed clear separation among the three clinical groups, reflecting distinct editing signatures (**Figure 6B**). Moreover, the total REB was markedly elevated in the MDS-H group compared to both MDS-L and MDS-C (**Figure S4C**).

**Figure 6.**
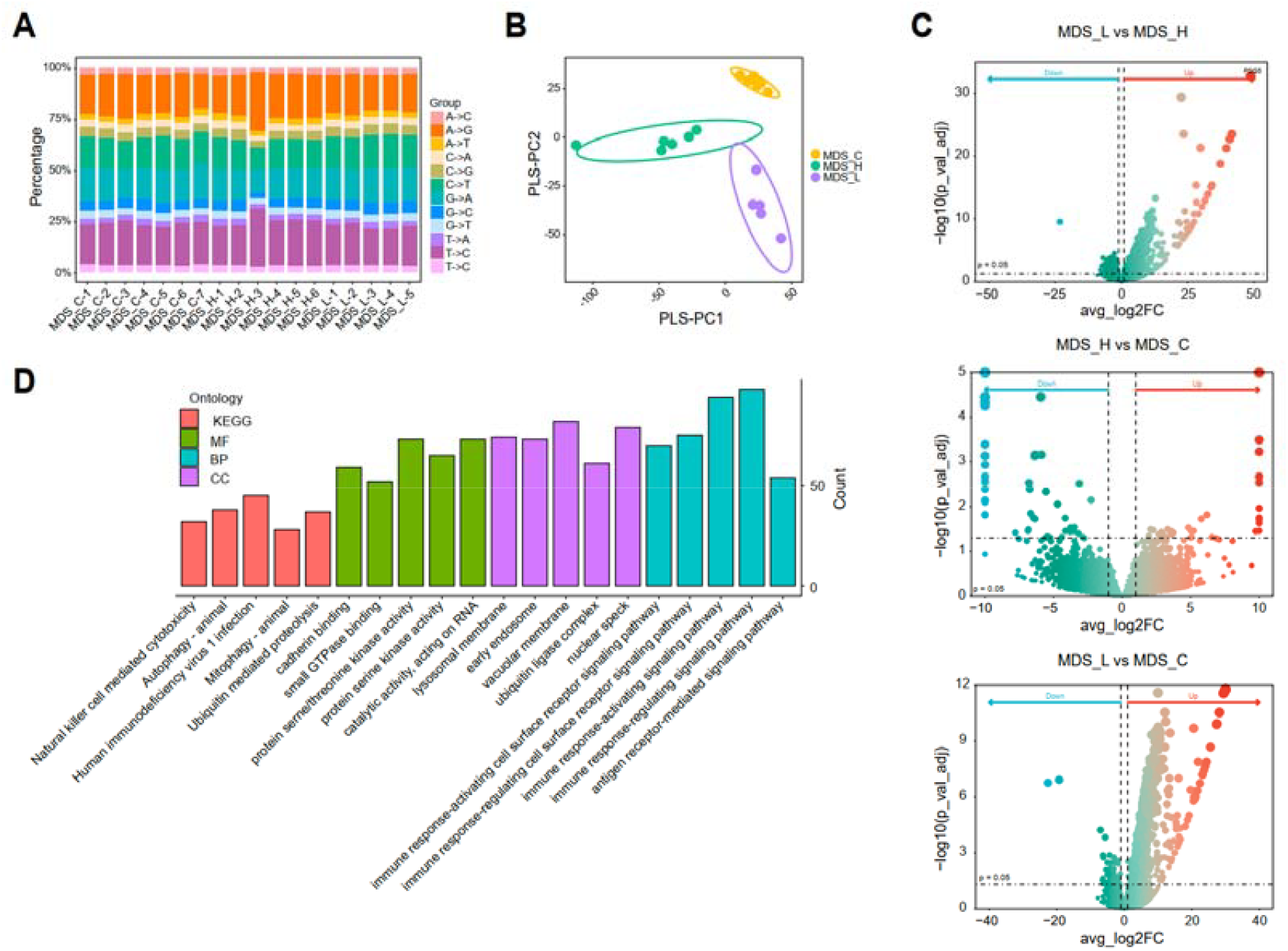
The landscape of RNA editing in MDS. (A) Distribution of RNA editing types across MDS-C, MDS-H, and MDS-L groups. (D) PCA of RNA editing-like events based on GERC metric. (G)Volcano plot of differentially edited genes using GERC metric. (H) GO enrichment of differentially edited genes.

Differential editing analysis revealed multiple genes with significantly altered editing levels across three groups (**Figure S4D**). A volcano plot (**Figure 6C**) highlighted high-confidence differentially edited genes. Enrichment analyses showed that these genes were significantly involved in immune signaling, apoptosis, and inflammatory pathways (**Figure 6D, Figure S4E**), with notable enrichment in NF-κB, IL-17, and TNF signaling cascades. Among these, PAICS, a key enzyme in purine biosynthesis, was significantly edited and upregulated in the MDS-H group, implicating it as a potential contributor to MDS progression.

### RNA editing in the dynamic drug treatment of melanoma

We next applied the RE-seq framework to study dynamic RNA editing changes during drug treatment in melanoma cells. Samples included untreated control cells (HDF) and cells exposed to a targeted therapy (PLX) for 4, 8, or 18 hours. RNA editing was quantified using the RELB metric at the locus level.

Canonical A-to-I and C-to-U events dominated the editing profile, which are mainly distributed in 3’UTR, exonic, and intronic regions (**Figure 7A, Figure S5A**). Editing consequence analysis revealed a majority of synonymous variants, though nonsynonymous and stop-gain edits were also observed (**Figure S5B**). PCA showed temporal separation among treatment groups, while total REB remained largely unchanged (**Figure 7B, Figure S5C**).

**Figure 7.**
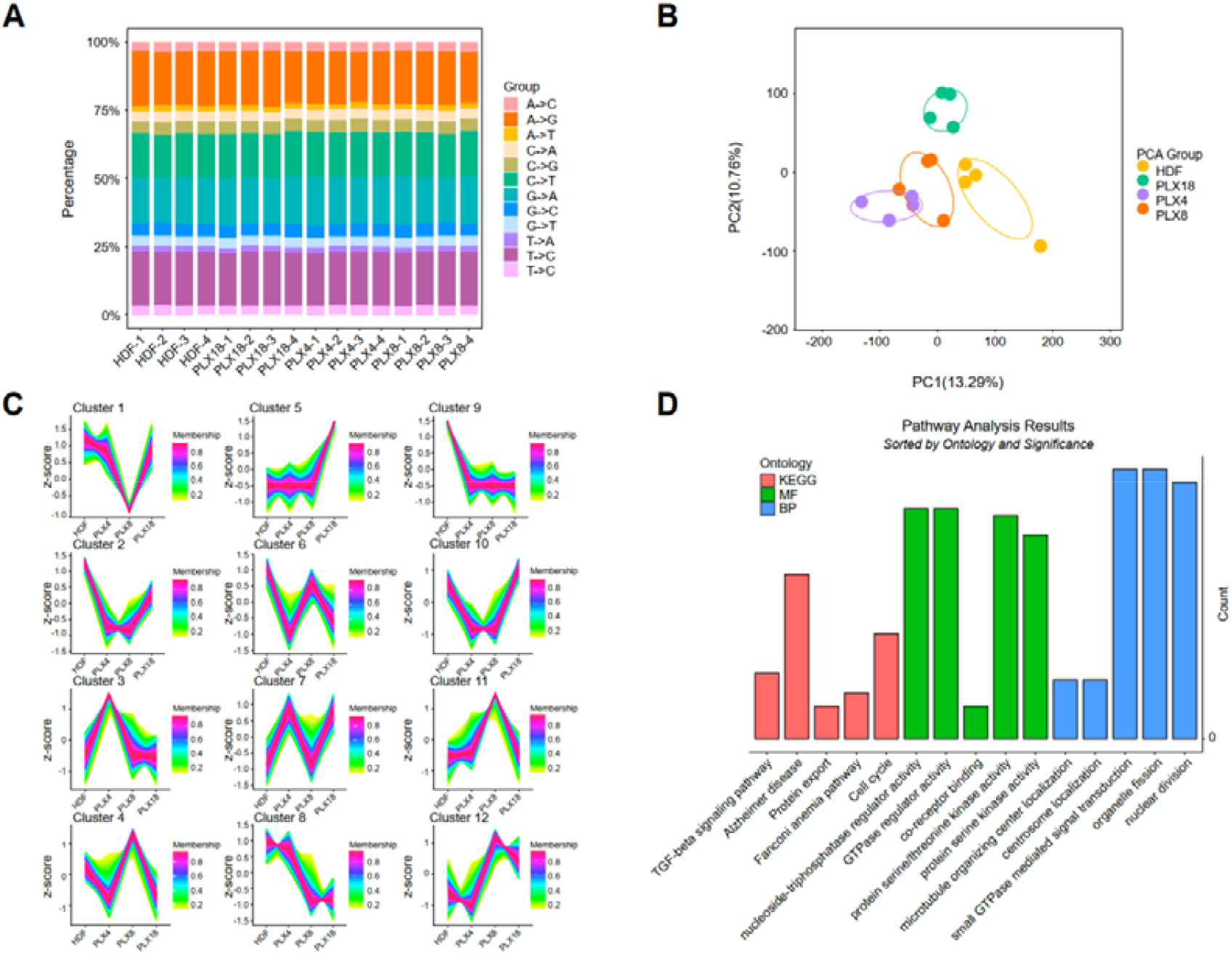
Dynamic RNA editing changes in melanoma drug treatment. (A) Distribution of RNA editing types across control and treated melanoma samples. (B) PCA of samples based on RELB metric. (C) Time-series distribution of dynamically regulated RNA editing-like loci. (D) GO enrichment of dynamically edited loci.

Time-course differential analysis using RELB identified dynamically regulated editing loci. Temporal editing trajectories revealed distinct editing programs induced by drug exposure (**Figure 7C**). Enrichment analysis of dynamically edited genes indicated involvement in nuclear division, mitotic cell cycle, small GTPase signaling, and TGF-β and mTOR signaling pathways. GSEA further highlighted the activation of HIF-1 signaling and metabolic reprogramming during drug response (**Figure 7D, Figure S5D**).

### RNA editing in colorectal cancer for human tissue

To explore RNA editing in colorectal cancer (CRC), we applied the RE-seq workflow to tumor and matched adjacent normal tissues (n = 10 pairs). RNA editing-like events were quantified at the locus level using LERC.

As in other human datasets, A-to-I and C-to-U events were dominant, primarily localized in intronic regions (**Figure 8A, Figure S6A**). Synonymous and nonsynonymous substitutions accounted for the majority of annotated editing consequences (**Figure S6B**). PCA analysis based on LERC values showed partial separation between tumor and normal samples (**Figure 8B**). Although REB did not differ significantly (p = 0.09), a trend toward elevated editing activity in tumors was noted (**Figure S6C**).

**Figure 8.**
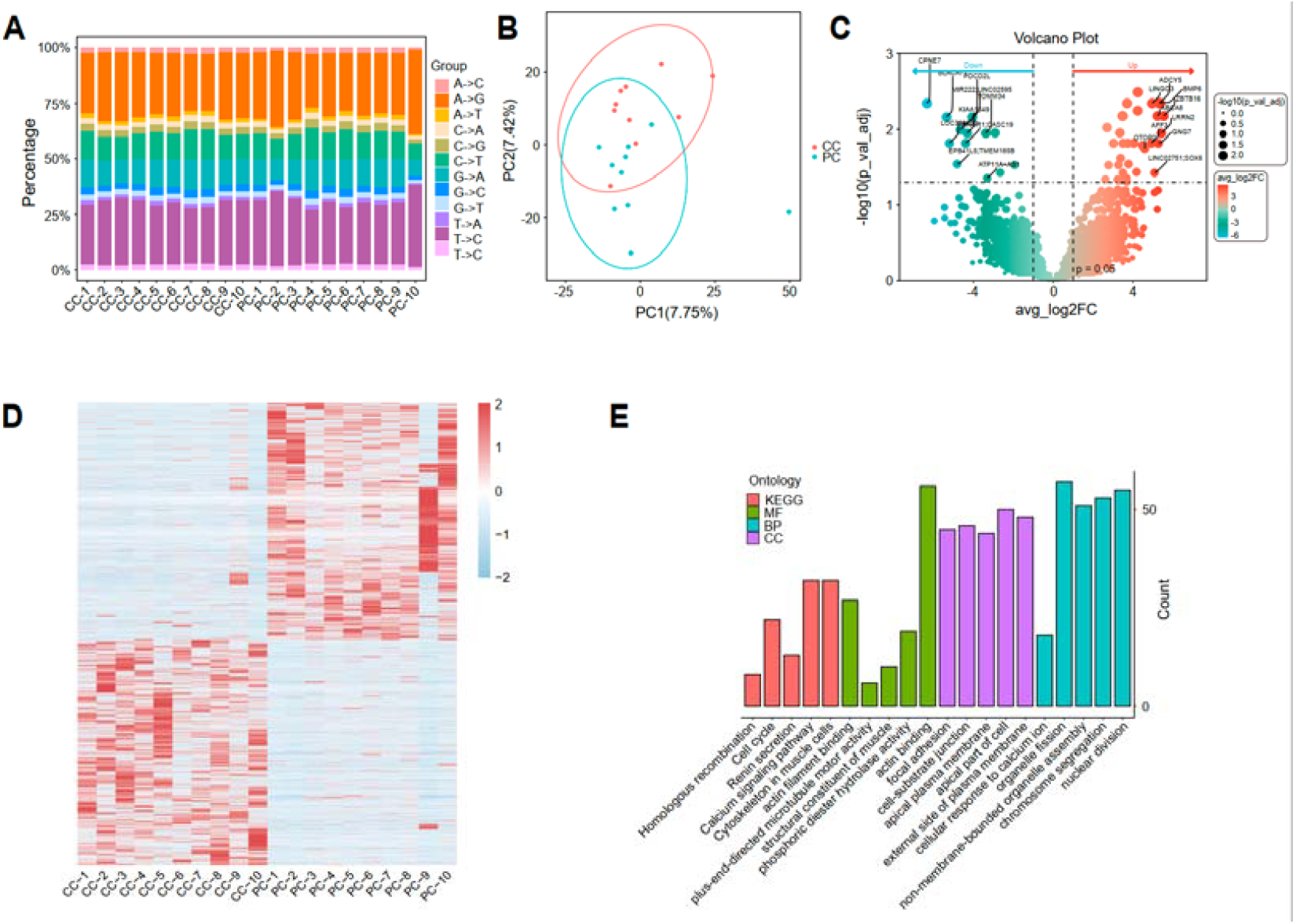
RNA editing profiles in human CRC. (A) Distribution of RNA editing types across CRC tumor and normal tissues. (B) PCA of samples based on LERC metric. (C) Volcano plot of differentially edited loci using LERC metric. (D) Heatmap of differentially edited loci using LERC metric. (E) GO enrichment of differentially edited genes.

Differential editing analysis using LERC identified site-specific alterations in RNA editing between tumor and normal tissues. A volcano plot of differentially edited loci is shown in **Figure 8C**, and a heatmap visualizing editing levels across samples is presented in **Figure 8D**. Functional enrichment analysis of genes linked to differentially edited loci suggested associations with immune regulation, cell cycle control, and protein processing in the endoplasmic reticulum (**Figure 8E, Figure S6D**), highlighting the potential functional relevance of RNA editing in CRC pathogenesis.

### Single-cell RNA editing landscape in colorectal cancer

To explore the cell-type-specific RNA editing landscape in colorectal cancer (COAD), we applied the scRE-seq pipeline to single-cell transcriptomic data from six tumor and matched normal tissue samples. Clustering based on single-cell RNA expression profiles identified ten major cell types without apparent batch effects (**Figure 9A–B**). To overcome the low per-cell coverage of editing events inherent to scRNA-seq, we employed a Spot-based strategy that aggregates 20 phenotypically similar cells per unit for editing quantification. Dimensionality reduction and clustering based on RNA editing features revealed inconsistencies with initial expression-based annotations. After correction, six high-confidence cell types were retained, and the revised UMAP projections for RNA expression and editing profiles showed improved concordance (**Figure 9C–D**). Quantification of REB number across cell types revealed elevated activity in epithelial and plasma cells (**Figure 9E**). Detailed characterization of editing types, genomic localization, and consequence categories further highlighted distinct cell-type-specific editing patterns (**Figure 9F, Figure S7A-B**).

**Figure 9.**
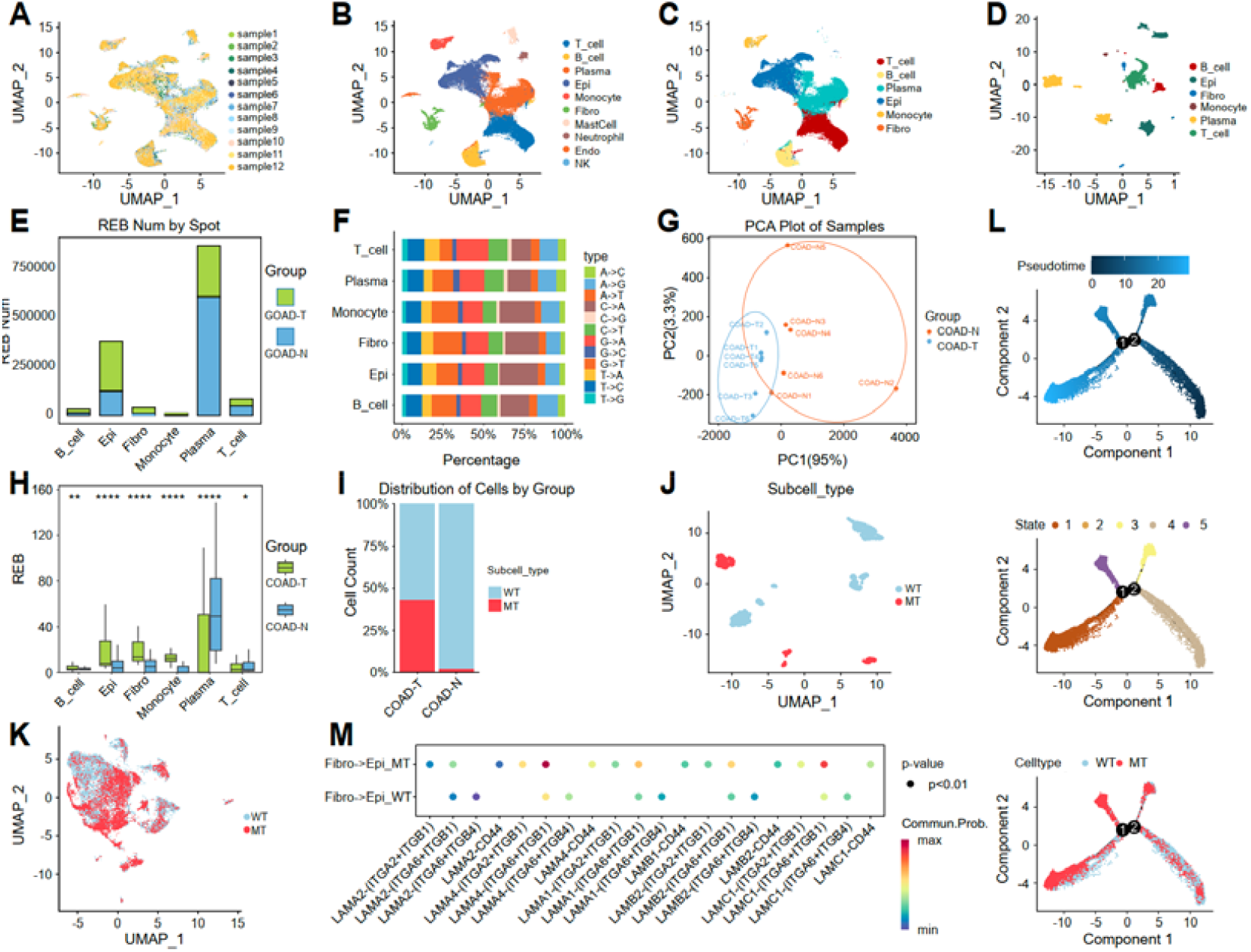
Single-cell RNA editing landscape in colorectal cancer. (A) UMAP clustering of cells based on gene expression profiles. (B) Cell type annotation using canonical markers (n = 10). (C) UMAP of corrected expression after annotation refinement. (D) UMAP clustering based on RNA editing features post Spot aggregation. (E) RNA editing burden across six major cell types. (F) Proportion of RNA editing types (A-to-I, C-to-U, etc.) across cell types. (G) PCA of RNA editing-like events in tumor (COAD-T) and normal (COAD-N) samples. (H) Comparison of REB across cell types in COAD-T vs COAD-N. (I) Proportions of wild-type (WT) and mutant (MT) epithelial cells across conditions. (J) UMAP clustering based on RNA editing status in epithelial cells. (K) UMAP clustering based on RNA expression in epithelial cells. (L) Pseudotime trajectory inference of epithelial subsets. (M) Cell–cell communication analysis of epithelial cells with surrounding cell types.

Principal component analysis of editing features clearly separated tumor (COAD-T) and normal (COAD-N) tissues, and cell-type-specific RNA editing markers were identified across the dataset (**Figure 9G, Figure S7C**). A comparison of REB between COAD-T and COAD-N samples indicated cell-type-dependent variation (**Figure 9H**). Focusing on epithelial cells, differential RNA editing analysis revealed several significantly altered sites between tumor and normal tissues (**Figure S7D**). Among these, an editing site in the FOS gene (chr14:75279987 C>T) was of particular interest. Based on the presence of this editing event, epithelial cells were stratified into wild-type (WT) and mutant (MT) groups. The proportions of WT and MT cells differed significantly between tumor and normal tissues (**Figure 9I**). Notably, RNA editing-based clustering effectively separated WT and MT cells, while expression-based clustering failed to do so, underscoring the added resolution offered by RNA editing (**Figure 9J-K**).

To explore the functional relevance of editing alterations, GO enrichment analysis of differentially edited genes in epithelial cells revealed enrichment in cytokine activity, chemokine receptor signaling, and metal ion sensing (**Figure S7E**). Single-cell trajectory inference further demonstrated distinct differentiation paths between WT and MT epithelial populations, accompanied by expression divergence and state-specific functional pathway enrichment (**Figure 9L, Figure S7F-H**). Finally, intercellular communication modeling identified altered interaction networks involving editing-positive epithelial cells, suggesting that RNA editing may shape tumor microenvironmental dynamics (**Figure 9M**).

### Integrated analysis of RE-seq and scRE-seq reveals RNA editing–mediated mechanisms underlying thymic atrophy recovery

To investigate the molecular mechanisms of *Ganoderma lucidum* spore–mediated recovery from thymic atrophy, we performed integrative bulk and single-cell RNA editing analyses using RE-seq and scRE-seq, respectively. Samples were collected from a murine thymic atrophy model induced by estradiol benzoate (Model), a control group (Control), and a treatment group receiving *Ganoderma lucidum* spores (GLS). Six single-cell RNA-seq datasets (two per group) and six matched bulk RNA-seq datasets (two per group) were processed to construct a multi-resolution RNA editing landscape (**Figure S8A**). Following annotation correction and quality control, we found that thymic samples were predominantly composed of T cells, which also exhibited substantial differences in REB across groups (**Figure 10A**). We then focused on T-cell subtypes to identify group-specific RNA editing patterns. Differential RNA editing analysis revealed several high-confidence sites distinguishing the Model group from both the Control and GLS groups (adjusted P < 0.05). Among these, the editing site chr11:62552979 A>G in the Ubb gene was consistently upregulated in the Model group and was selected as a key editing marker (**Figure 10B–C**). This candidate RNA editing site was used to stratify T cells into wild-type (WT) and mutant-type (MT) populations. Subset enrichment analysis revealed that MT cells were significantly enriched in membrane invagination-related pathways (**Figure S8B**), suggesting functional relevance to thymic involution.

**Figure 10.**
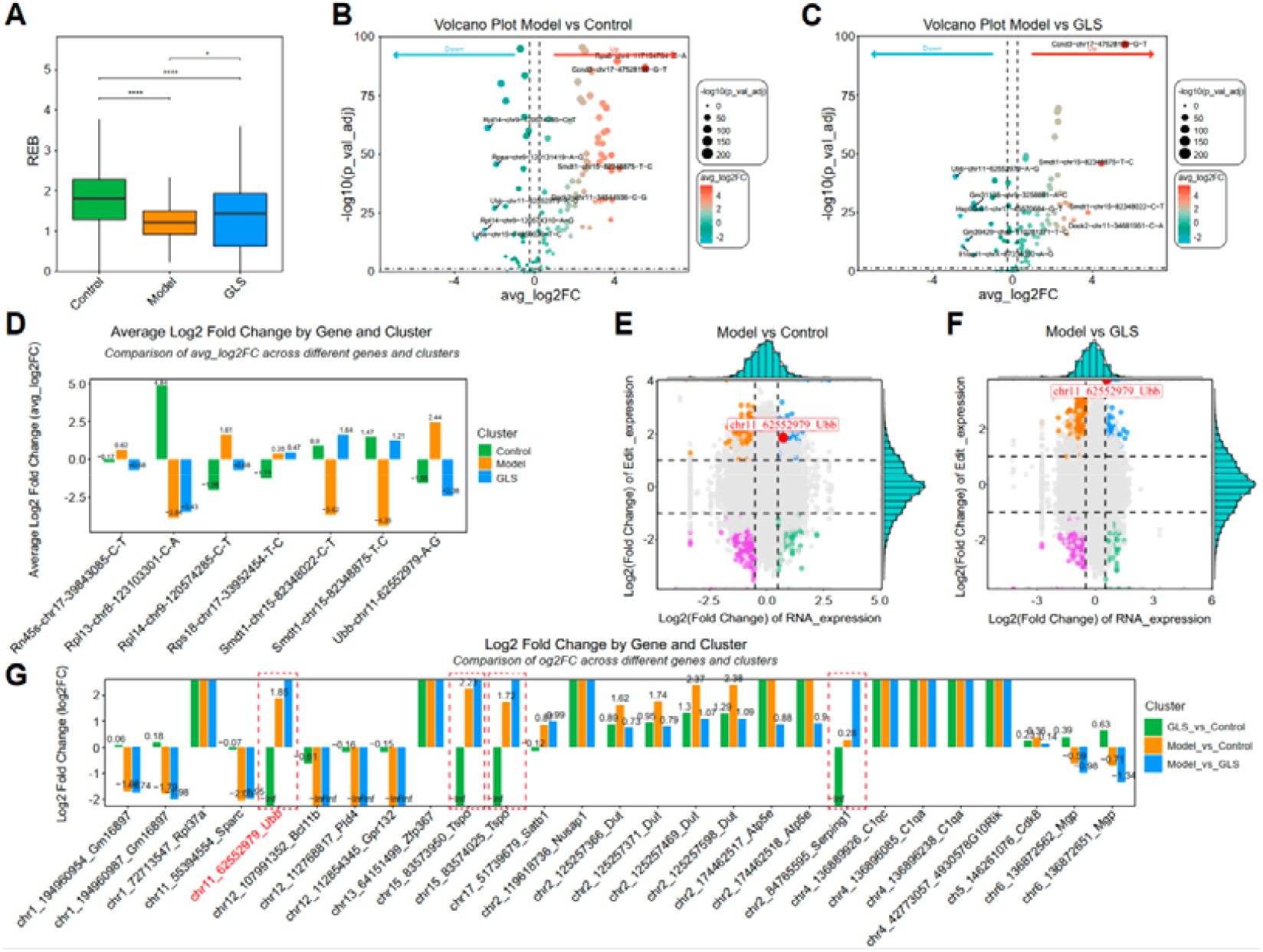
OmniRE-seq analysis of thymic atrophy and recovery in a mouse model. (A) Boxplot comparing REB in T cell subsets across Control, Model, and GLS groups. (B) Volcano plot of differential RNA editing sites between Model and Control in T cells (scRE-seq). (C) Volcano plot of differential RNA editing sites between Model and GLS in T cells (scRE-seq). (D) Shared differential RNA editing events in the MT subpopulation detected across all comparisons by scRE-seq. (E) Differentially edited sites across all three groups in bulk RE-seq data. (F) Nine-quadrant plot comparing RNA editing and expression for Model vs. Control in RE-seq data. (G) Nine-quadrant plot comparing RNA editing and expression for Model vs. GLS in RE-seq data.

We next identified editing events shared across all three experimental comparisons within the MT population, selecting only those detected by both scRE-seq and RE-seq pipelines. Two overlapping sites were identified: Ubb-chr11:62552979 A>G and Rpl14-chr9:120574285 C>T (**Figure 10D**). These showed a pattern of upregulation in the Model group and downregulation in both the Control and GLS groups, implicating them in thymic atrophy and its reversal. We employed quadrant-based joint analysis to evaluate editing-expression relationships. In both Model vs. Control and Model vs. GLS comparisons, Ubb exhibited consistent quadrant positioning, supporting its potential role as an RNA editing–mediated regulator in thymic remodeling (**Figure 10E–F**).

To validate these findings in bulk tissue, we applied RE-seq to the matched bulk RNA-seq samples. Consistent with the scRE-seq results, the Ubb editing site was also significantly altered in the bulk data, reinforcing its biological significance (**Figure 10G**). Moreover, we observed coordinated trends between editing levels and gene expression for Ubb in both Model vs. Control and Model vs. GLS comparisons, further highlighting its regulatory importance. Together, these findings demonstrate the power of multi-resolution editing analysis to uncover functionally relevant RNA editing events and highlight Ubb-chr11:62552979 A>G as a key editing signature involved in both thymic atrophy and its therapeutic reversal by *Ganoderma lucidum* spore intervention.

### Discussion

RNA editing is an essential post-transcriptional modification mechanism that introduces nucleotide substitutions in RNA molecules, playing pivotal roles in transcriptome diversity, cellular adaptation, and disease pathogenesis ^40^. Despite its biological significance, systematic and quantitative evaluation of RNA editing remains technically challenging, particularly due to limited coverage in RNA-seq, variable expression across conditions, and the lack of standardized analysis frameworks. In this study, we established OmniRE-seq, a dual-layered computational framework comprising RE-seq for bulk RNA-seq and scRE-seq for single-cell data, providing scalable and reproducible pipelines to decipher RNA editing dynamics across biological contexts.

The RE-seq pipeline advances RNA editing analysis in bulk transcriptomes by introducing a multi-level quantification system and matching distribution-specific statistical tools. Editing signals are quantified at the loci, gene and sample levels via three metrics: raw read counts (LERC/GERC), normalized editing frequency (RELB/REGB), and transcriptome-wide burden (REB). Count-based metrics are assessed using DESeq2 or limma, which model overdispersed counts via negative binomial or linear models. In contrast, burden-based metrics are not count-distributed and are analyzed via permutation-based frameworks to avoid parametric assumptions, especially in small cohorts.

The choice of quantification metric is context specific. In the sepsis model, global shifts in RNA editing patterns are evident from both PCA and REB analyses, prompting the use of REGB to capture gene-level burden changes across disease stages. In MDS analysis, strong separation in PCA but only partial REB divergence leads us to use GERC, focusing on raw gene-level editing activity. In contrast, melanoma and colorectal cancer datasets did not show marked REB shifts, and differential editing was better captured at the site level via RELB and LERC, respectively. These examples underscore that editing burden metrics offer normalization advantages for heterogeneous data, while read count-based metrics retain sensitivity in modest or localized editing changes.

Beyond quantification, RE-seq integrates functional enrichment and immune cell deconvolution, enabling insight into how RNA editing reshapes regulatory networks. The REB, analogous to Tumor Mutational Burden (TMB) ^41^, emerges as a promising biomarker in precision oncology. Its global nature facilitates patient stratification and may predict immunotherapy response, as elevated editing burden has been linked to neoantigen generation, immune evasion, or inflammation-associated pathologies. Future clinical validation of REB in immunotherapy cohorts could expand its utility as a transcriptome-derived immunogenomic feature ^42^.

To resolve cellular heterogeneity in RNA editing, we developed scRE-seq, which applies the RE-seq logic to 10x Genomics scRNA-seq datasets. Recognizing the sparsity of single-cell transcriptomes, scRE-seq introduces the “Spot” strategy: pooling 20 cells of the same individual and cell type to generate pseudo-bulk units. This approach improves read depth while retaining resolution sufficient for cell-type-specific quantification. Furthermore, integration of scRNA-derived annotations with editing data allows rigorous quality control, eliminating clusters with high annotation discordance or editing inconsistency. This correction enhances specificity and permits detection of editing heterogeneity across immune, epithelial, and stromal cell types.

The power of scRE-seq lies in its ability to decouple RNA editing variation from gene expression. As demonstrated in colorectal cancer, epithelial subclusters defined by differential editing of the FOS gene could not be resolved by transcriptomic data alone. Editing-defined subpopulations showed divergent differentiation trajectories and interaction networks, revealing that RNA editing can reprogram cell states independently of transcriptional output. These findings support scRE-seq as a critical addition to single-cell analysis, particularly in cancer and immunology, where post-transcriptional regulation is key.

The integration of RE-seq and scRE-seq in a mouse model of thymic atrophy treated with *Ganoderma lucidum* spores further underscores the potential of multi-resolution RNA editing analysis. Both pipelines converged on a shared editing site (Ubb-chr11-62552979-A-G) implicated in T cell subpopulation dysfunction. Such convergence from bulk and single-cell data not only validates candidate events but also highlights their cellular context and potential functional implications. This suggests that OmniRE-seq enables a more complete mechanistic understanding by bridging tissue-level signals with cellular specificity.

It is important to clarify that RNA editing–mediated nucleotide substitutions differ fundamentally from DNA mutations in both mechanism and biological implication. Most therapeutic agents are not designed to directly alter RNA sequences at specific sites. However, environmental changes or pharmacological interventions can indirectly modulate RNA editing activity by affecting the expression or enzymatic activity of ADAR/APOBEC family members, altering RNA secondary structure, or changing substrate accessibility. These indirect effects may shift editing frequencies at certain loci, particularly in non-coding regions, without introducing permanent changes to the genome. In our melanoma time-course dataset, we observed sustained yet reversible alterations in RNA editing profiles following drug exposure, illustrating that editing patterns can reflect dynamic cellular responses rather than irreversible genetic lesions. Given that RNA editing is inherently heterogeneous across individuals, tissues, and physiological states, and that the majority of observed edits occur in untranslated or intronic regions, such changes are more plausibly interpreted as part of the transcriptomic regulatory landscape than as genotoxic events. This distinction is critical for safety considerations. While DNA-level base substitutions can pose long-term mutagenic risks, RNA editing–related sequence changes are transient, reversible, and integrated into normal adaptive responses.

Nonetheless, challenges remain in interpreting the functional consequences of RNA editing. Many events occur in non-coding or intronic regions, and even protein-coding edits often lack clear phenotypic annotation. While our study employed GO, KEGG, and GSEA for downstream analysis, deeper mechanistic insights will require integration with proteomics ^43^, ribosome profiling, or spatial transcriptomics. Moreover, distinguishing editing from SNPs, accounting for isoform diversity, and integrating genotype–phenotype links (e.g., edQTLs) remain important future directions. Future studies should further evaluate context-dependent editing changes under therapeutic exposure, with careful distinction between dynamic transcriptomic regulation and permanent genomic alterations.

The framework presented here lays a foundation for translational and biotechnological applications beyond oncology ^44^. RNA editing is emerging as a regulatory mechanism in plant stress responses ^45^, algal metabolism ^46^, and immune development ^47^, with potential for metabolic engineering and cell fate modulation ^48^. Our broader application of editing analysis in microalgae and herbal intervention models (e.g., *Ganoderma lucidum* spore recovery from thymic atrophy) reflects the versatility of OmniRE-seq across species, tissues, and perturbations.

In conclusion, OmniRE-seq unifies bulk and single-cell RNA editing analysis into a standardized, modular, and scalable framework. Its dual pipelines, RE-seq and scRE-seq, enable high-resolution quantification, cross-condition comparison, and mechanistic interpretation of RNA editing dynamics. By facilitating robust identification of editing biomarkers and functional candidates, OmniRE-seq provides a powerful platform for advancing RNA editing research and unlocking its potential in diagnostics, therapeutics, and systems biology.

## Methods

### Bulk RE-seq analysis

The Bulk RE-seq pipeline processes RNA-seq raw FASTQ data through a series of computational steps, including quality control, alignment, variant calling, and differential analysis. This pipeline is designed to systematically identify and quantify RNA editing events, enabling the discovery of disease-related mechanisms.

Raw sequencing data (FASTQ files) were subjected to quality control using Fastp v0.23.2 ^30^ to remove low-quality reads and contaminants. Cleaned reads were aligned to the reference genome using HISAT2 v2.2.1 ^31^, and the resulting BAM files were sorted using SAMtools v1.10.2 ^32^. PCR duplicates were removed using Picard (http://broadinstitute.github.io/picard). Candidate RNA editing sites were initially called using GATK v4.2.0, and subsequently fitered to remove known SNPs annotated in dbSNP/known SNP databases, low depth sites and low-quality sites ^34^. The retainted high-confidence sites were annotated using ANNOVAR ^35^, and the distribution of RNA editing-like events across genomic regions was analyzed.

RNA editing quantification was performed at three levels: loci-level, gene-level, and sample-level. For loci-level quantification, we defined the following metrics:

#### Loci Read Counts (LRC)

The number of reads supporting an mapped to the site at a specific genomic locus.

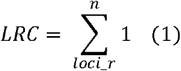

*Loci_r*: Number of reads/fragments at a specific site

#### Loci Editing Read Counts (LERC)

The number of reads supporting an RNA editing-like event at a specific genomic locus.

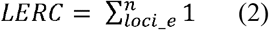

*Loci_r*: Number of edited reads/fragments at a specific site

#### Loci Editing Frequency (LEF)

The proportion of edited reads relative to the total reads mapped to a specific site.

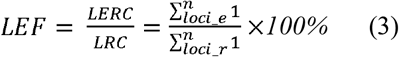

#### RNA Editing Loci Burden (RELB)

A normalized measure of RNA editing activity at a specific site, accounting for gene length and sequencing depth.

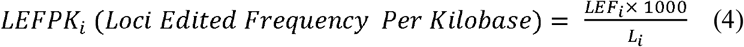

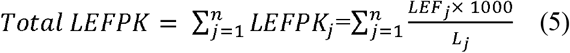

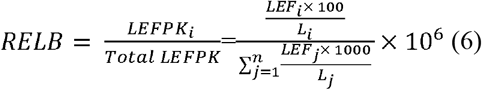

*L*_*i*_: The total exon length of gene i (unit: base pairs, bp). *n*:The total number of genes in the sample.

At the gene level, we quantified RNA editing activity across all editing sites within a gene:

#### Gene Read Counts (GRC)

The number of reads supporting an mapped to the gene at a specific genomic locus.

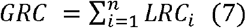

where *n* is the number of editing sites within the gene.

#### Gene Editing Read Counts (GERC)

The total number of edited reads across all editing sites within a gene.

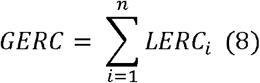

where *n* is the number of editing sites within the gene.

#### Gene Editing Frequency (GEF)

The proportion of edited reads relative to the total reads mapped to the gene.

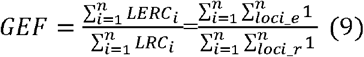

#### RNA Editing Gene Burden (REGB)

A normalized measure of RNA editing activity at the gene level, accounting for gene length and sequencing depth.

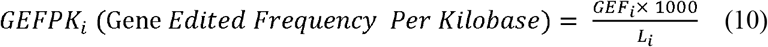

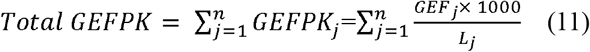

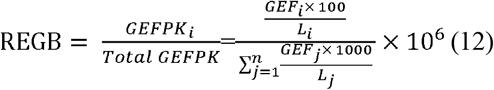

*N*_*i*_: The proportion of gene reads relative to the total reads mapped to a specific gene. *L*_i_: The total exon length of gene i (unit: base pairs, bp). *n*:The total number of genes in the sample.

At the sample level, we quantified the overall RNA editing activity across all genes in the transcriptome:

#### RNA Editing Burden (REB)

REB number normalized by the total protein-coding sequence length in the transcriptome.

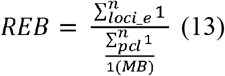

*pcl*: Protein coding length (bp).

Differential RNA editing analysis was performed using DESeq2 v1.42.1 ^36^ for count-based metrics (LERC and GERC) and Permutation-based methods for burden-based metrics (RELB and REGB). The permutation approach was chosen due to its ability to handle small sample sizes and non-normally distributed data. Traditional methods like t-test and DESeq2 assume normality, which often leads to inaccurate results in RNA editing datasets. Permutation testing, on the other hand, does not rely on distributional assumptions and provides more reliable p-values by shuffling the data and recalculating the test statistic multiple times to generate a null distribution.

The formula for permutation-based p-value calculation is:

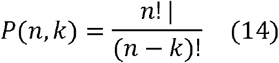

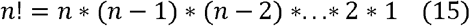

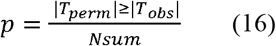

Let *n* denote the total number of elements in set S, and *k* denote the number of elements selected from set S. *n*! denotes the factorial of *n*, which is the product of all positive integers up to *n*. where *T*_*obs*_ is the observed test statistic, and *T*_*perm*_ is the test statistic from permuted data. *Nsum:* Total number of permutations

The quantification of RNA editing-like events at the loci, gene, and sample levels provides a comprehensive framework for analyzing RNA editing activity. The use of normalized metrics (e.g., RELB, REGB, REB) ensures that comparisons across different conditions and samples are robust and unbiased. The permutation-based approach further enhances the reliability of the results, especially in cases where traditional statistical methods (e.g., t-test, DESeq2) may fail due to small sample sizes or non-normal data distributions.

Functional enrichment analysis was performed using the clusterProfiler. Gene Ontology (GO) terms and Kyoto Encyclopedia of Genes and Genomes (KEGG) pathways were analyzed to identify biological processes, cellular components, and molecular functions associated with differentially edited genes. The enrichment analysis was conducted with the following parameters: *pvalueCutoff* = 0.05, *pAdjustMethod* = BH, and *qvalueCutoff* = 0.5.

All visualizations and statistical analyses were performed using R v4.3.3 (https://www.r-project.org/) and the Bioconductor (https://bioconductor.org/) software library. Differential editing events were visualized using volcano plots, and functional enrichment results were displayed using bubble charts and bar plots.

### Single cell RE-seq analysis

To investigate RNA editing dynamics at single-cell resolution, we developed and applied the scRE-seq pipeline, optimized for 10x Genomics Chromium single-cell RNA-seq data. The pipeline is designed to address challenges such as low transcript coverage and technical sparsity inherent to droplet-based single-cell platforms by introducing a pseudo-bulk “Spot”-based strategy for robust editing detection.

Raw FASTQ files were processed using the CellRanger pipeline (v6.1.2) with reference genomes GRCh38 (human) or MM10 (mouse), generating aligned BAM files and gene-cell expression matrices. Downstream analysis was performed using the Seurat R package (v4.3.1). Low-quality cells were filtered based on the following criteria: (1) cells expressing 200–10,000 genes; (2) mitochondrial gene content <30%; (3) hemoglobin gene expression <5%; and (4) dissociation-related signature scores within the top 95% of all cells. Data were normalized using the NormalizeData function, and highly variable genes were identified via FindVariableFeatures using the vst method.

Dimensionality reduction was performed using principal component analysis (PCA), and the top 30 principal components were selected for Uniform Manifold Approximation and Projection (UMAP) visualization. Batch effects across samples were corrected using

Harmony. Clustering was performed using shared nearest neighbor (SNN) graph construction (FindNeighbors) and community detection (FindClusters) with resolution parameters optimized for biological interpretability. Cell type annotation was achieved using a curated marker gene reference and validated by PCA, UMAP, and heatmap visualization.

To enhance RNA editing detection, we constructed Spots by aggregating 20 transcriptomically similar cells from the same individual and annotated cell type. Spot-specific BAM files were extracted using cell barcodes (CBs), and PCR duplicates were stringently removed based on both CB and unique molecular identifiers (UMIs). RNA editing-like sites were detected using REDItools, a high-sensitivity toolkit for RNA editing identification in low-coverage RNA-seq data ^49^. Identified editing events were aggregated into a spot-level editing matrix.

The formula for RNA editing frequency in a Spot (SREF) is:

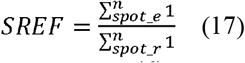

*spot_e*: Total edited reads/fragments at a specific spot. *spot_r*: Total reads/fragments at a specific spot.

Dimensionality reduction and clustering were also applied to the RNA editing matrix, using the same PCA and UMAP strategies as for gene expression data. Multimodal refinement was implemented by cross-referencing editing-based and expression-based clusters, and annotation-inconsistent clusters (e.g., editing clusters with >40% mixed cell types) were excluded to enhance annotation accuracy.

Differential RNA editing events were identified using group-wise comparisons within each cell type. Volcano plots and heatmaps were generated to visualize editing dynamics. Cell-type-specific RNA editing burden (REB) was computed to assess editing intensity across conditions.

Genes harboring differentially edited sites were subjected to Gene Ontology (GO) and Kyoto Encyclopedia of Genes and Genomes (KEGG) enrichment analyses using the ClusterProfiler package. Adjusted p-values were calculated using the Benjamini–Hochberg (BH) method with thresholds of p.adjust < 0.05 and qvalue < 0.5.

Pseudotemporal trajectories were inferred using Monocle 2 to reconstruct differentiation dynamics of specific cell populations. Genes associated with trajectory branches were analyzed for functional enrichment to identify stage-specific regulatory programs.

Intercellular signaling networks were inferred using CellChat (v1.6.0), which integrates ligand-receptor pairs and cofactor interactions. Expression data were normalized by cluster, and interaction probabilities were estimated across cell types. Interaction networks were visualized to reveal alterations in communication strength and pathway specificity under different biological conditions.

## Data Availability

The RE-seq case study datasets were obtained from multiple sources. Mouse sepsis RNA-seq data were provided by Professor Shengbao Wang and are available from the corresponding author upon reasonable request. Myelodysplastic syndrome (MDS) datasets were provided by Professor Xiao Yan and are likewise available upon request. The melanoma RNA-seq data were retrieved from the public repository under BioProject accession PRJNA779028, and the colorectal cancer bulk RNA-seq data are publicly available under BioProject accession PRJNA1047096. For the scRE-seq analysis, single-cell RNA-seq datasets of colorectal cancer were provided by Professor Yibin Zhao and are available from the corresponding author upon reasonable request. The mouse thymic atrophy model treated with *Ganoderma lucidum* spores was provided by Dr. Jihong Yang, and access can be granted upon reasonable request. The complete analysis pipelines for RE-seq and scRE-seq are available at https://github.com/biotranstech/RE-Seq and https://github.com/biotranstech/scRNA-Seq respectively.

## Author’s contributions

Y.Z. and Y.H. designed and supervised the project. W. L., P.L., and J.Y. performed the analysis and wrote the code. Y.H. and J.Y. wrote the paper. All authors read, edited and approved the final manuscript.

## Acknowledgements

We thank Shengbao Wang, Xiao Yan, and Yibin Zhao for providing us the test datasets used in the case studies and evaluation of the OmniRE-seq framework.

## Supplementary Figures

**Supplementary Figure 1.**
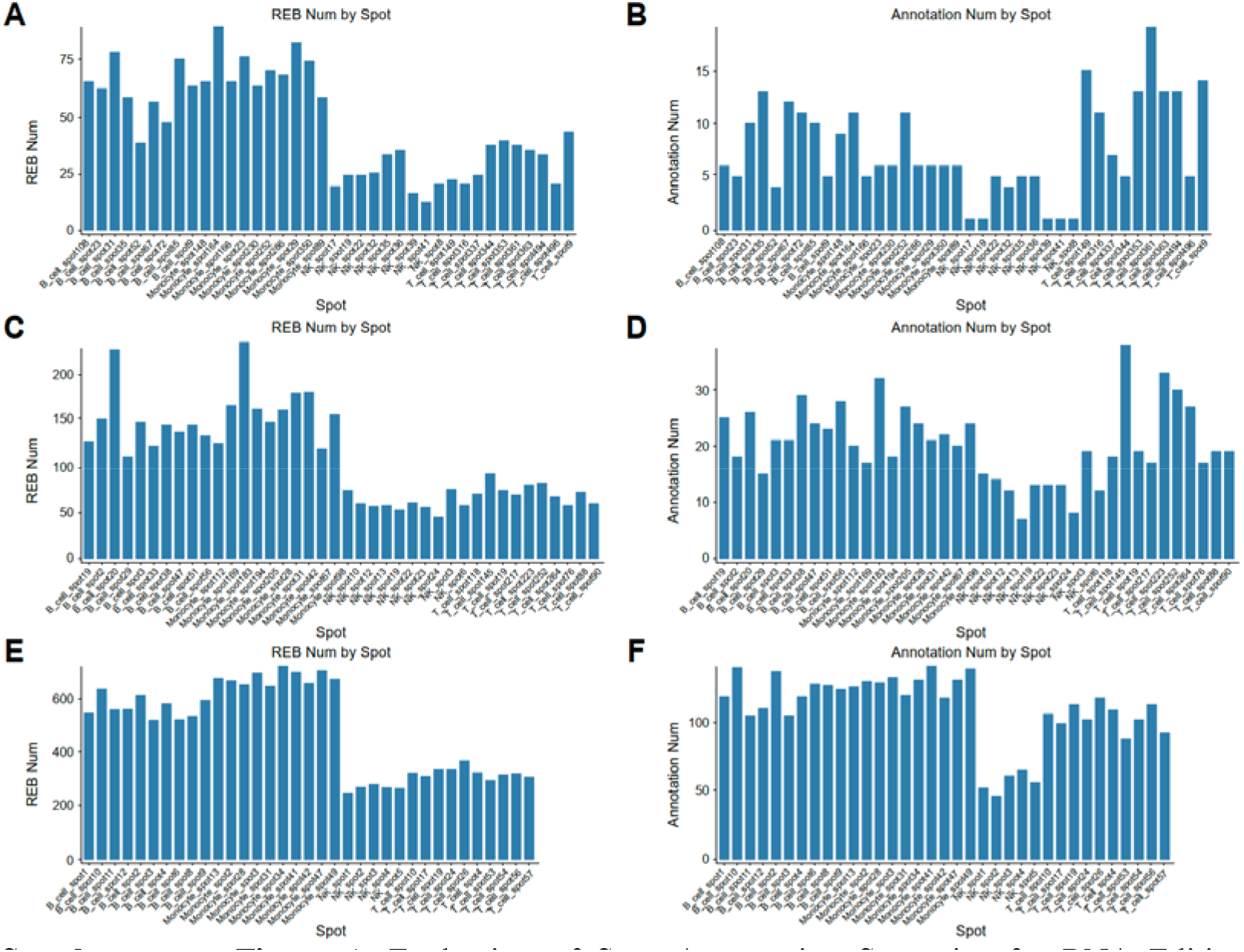
Evaluation of Spot Aggregation Strategies for RNA Editing Detection in scRE-seq. (A) REB Number identified using a Spot size of 10 cells. (B) Functionally annotated REB Number identified using a Spot size of 10 cells. (C) REB Number identified using a Spot size of 20 cells. (D) Functionally annotated REB Number identified using a Spot size of 20 cells. (E) REB Number identified using a Spot size of 90 cells. (F) Functionally annotated REB Number identified using a Spot size of 90 cells.

**Supplementary Figure 2.**
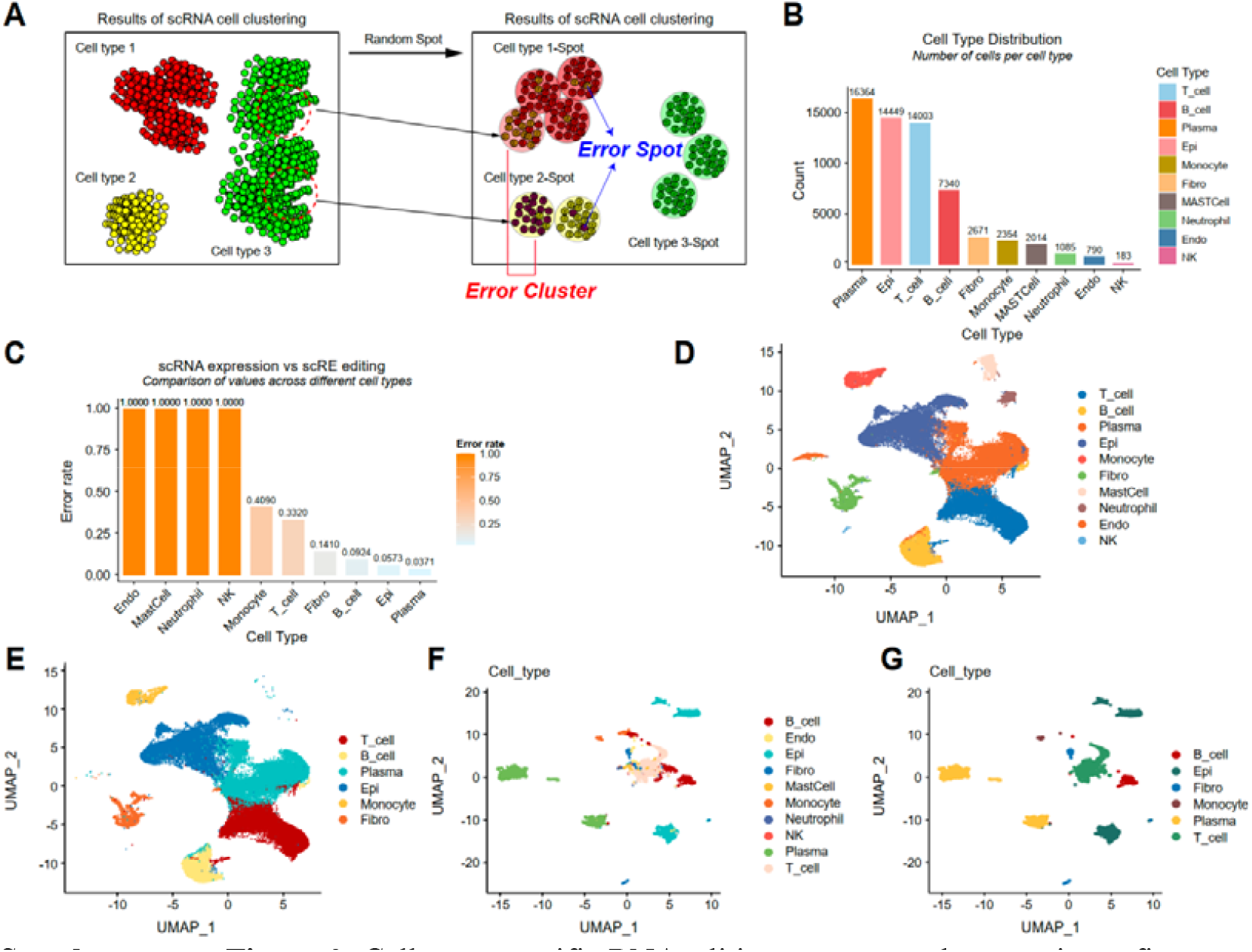
Cell-type-specific RNA editing patterns and annotation refinement. (A) Error cluster filtering principle. (B) Number of cells before correction. (C) Cell type error rate. (D) Single cell expression cluster results before correction. (E) Correction of pre-single-cell RNA editing clustering results. (F) Single cell expression cluster map after correction. (G) Single cell RNA edited cluster map after correction.

**Supplementary Figure 3.**
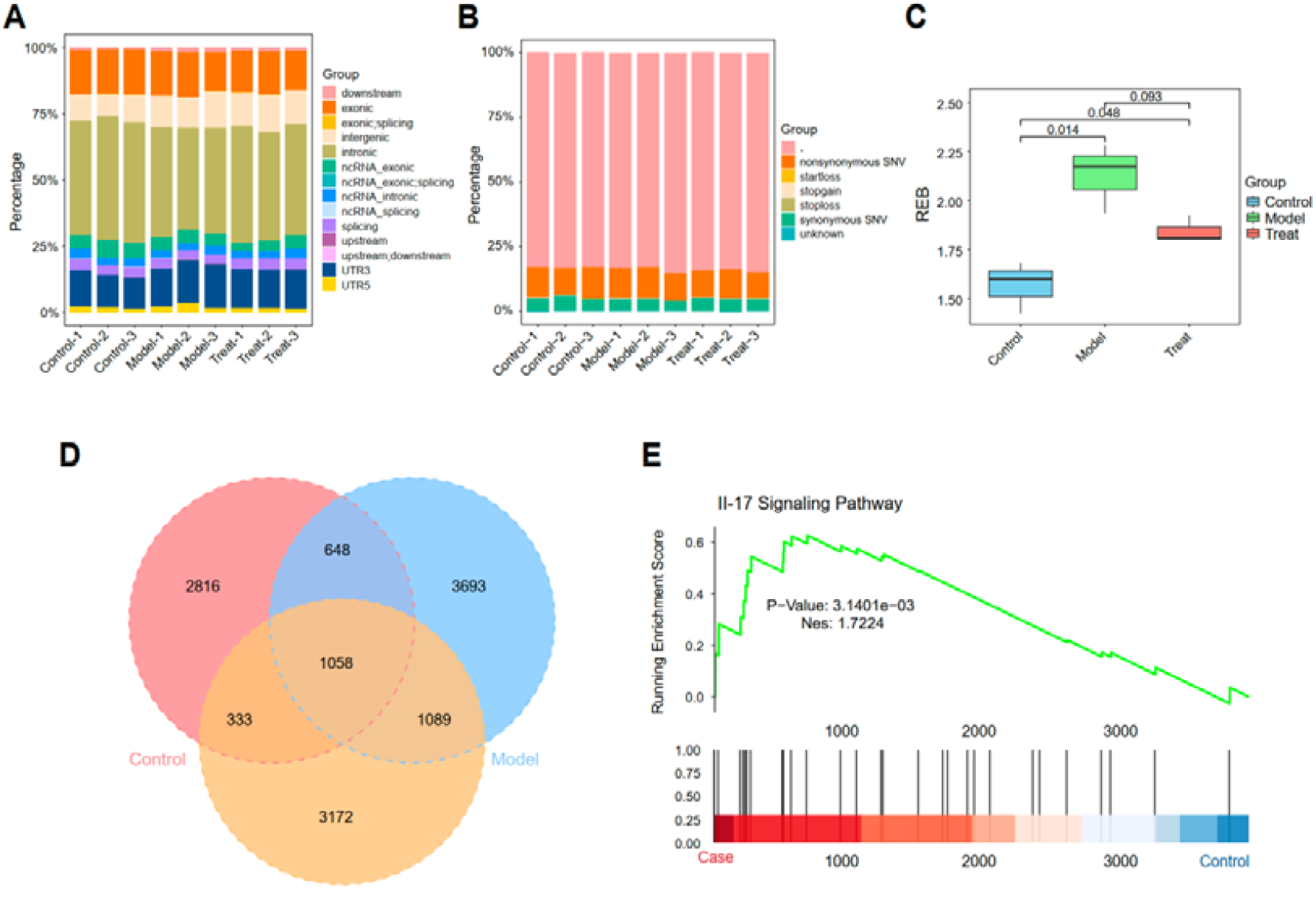
Extended characterization of RNA editing in sepsis-induced cardiac injury. (A) Genomic region distribution of RNA editing-like sites. (B) Distribution of RNA editing functional consequences. (C) REB comparison across the three groups. (D) Venn diagram of differentially edited genes across groups. (E) GSEA enrichment results highlighting relevant pathways.

**Supplementary Figure 4.**
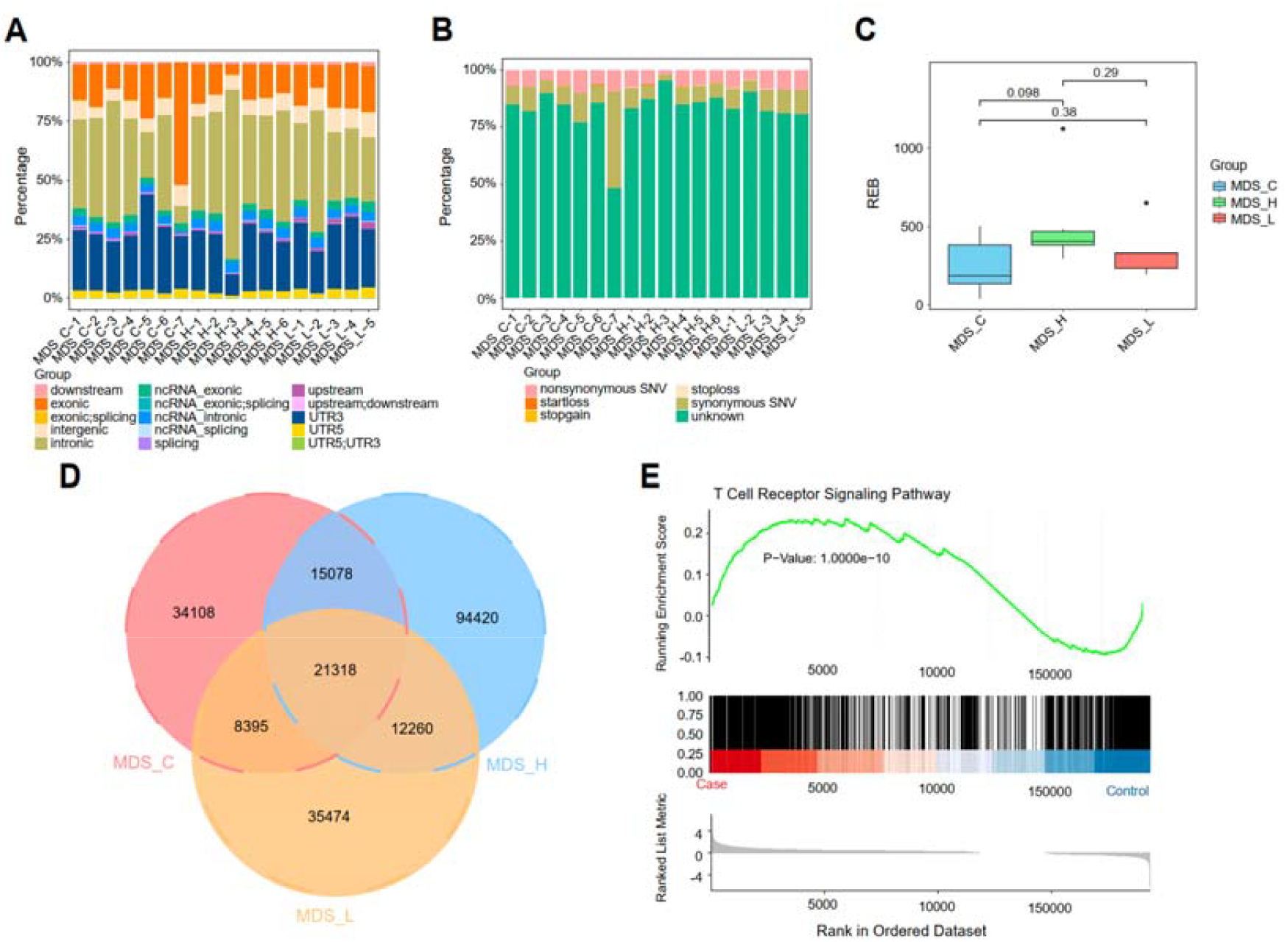
Extended RNA editing analysis in MDS. (A) Genomic region annotation of RNA editing-like events. (B) Functional classification of RNA editing consequences. (C) REB comparison across MDS groups. (D) Venn diagram of differentially edited genes across groups. (E) GSEA enrichment associated with differential editing.

**Supplementary Figure 5.**
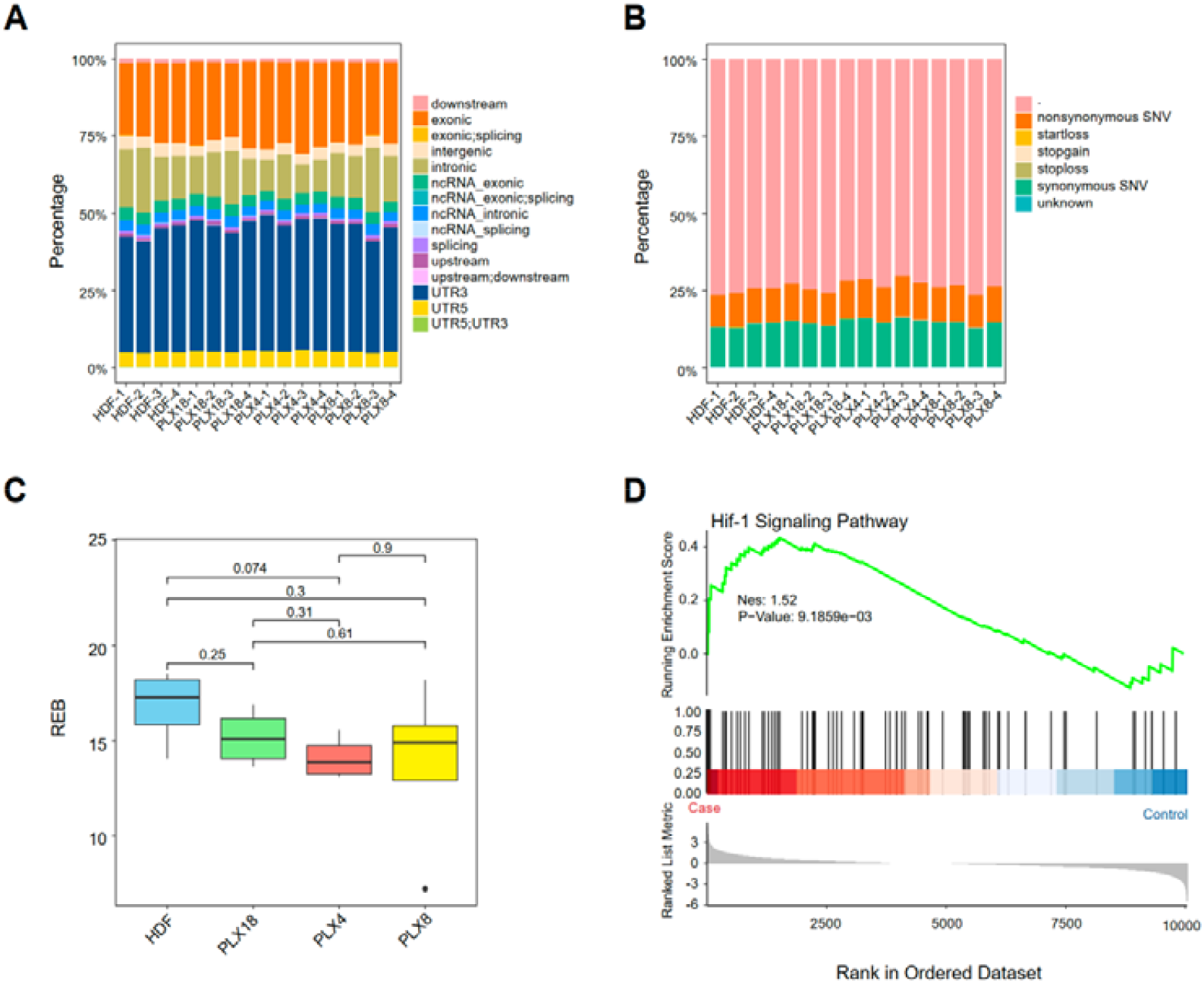
Extended RNA editing characterization of melanoma drug treatment analysis. (A) Genomic region classification of RNA editing-like loci. (B) Functional consequences of identified editing events. (C) REB comparison between HDF and drug-treated samples (PLX4, PLX8, and PLX18). (D) GSEA enrichment associated with differential editing events.

**Supplementary Figure 6.**
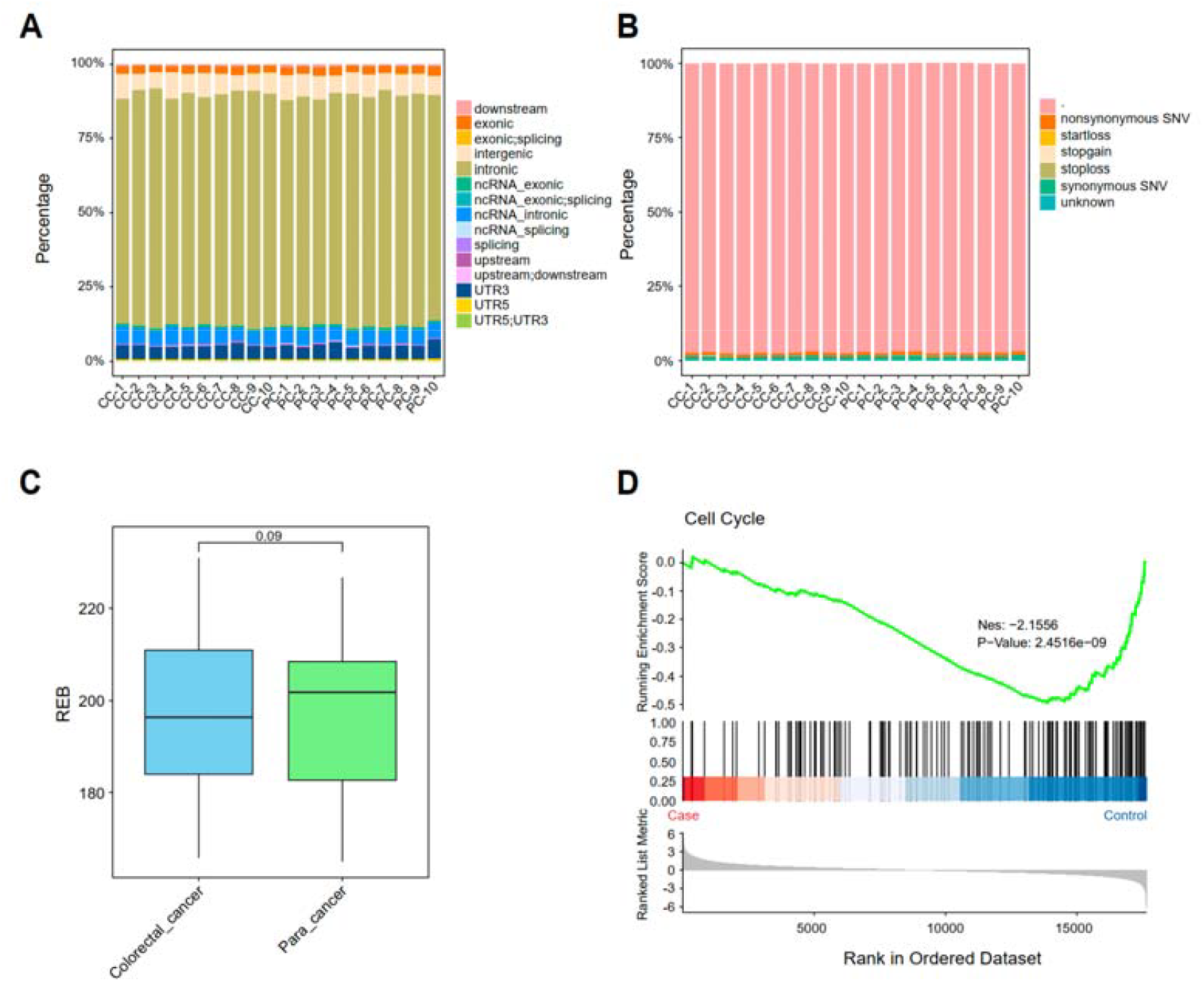
Extended RNA editing characterization of human CRC. (A) Genomic region annotation of candidate RNA editing-like sites. (B) Functional classification of RNA editing consequences. (C) REB comparison across tumor and normal tissues. (D) GSEA enrichment associated with differential editing events.

**Supplementary Figure 7.**
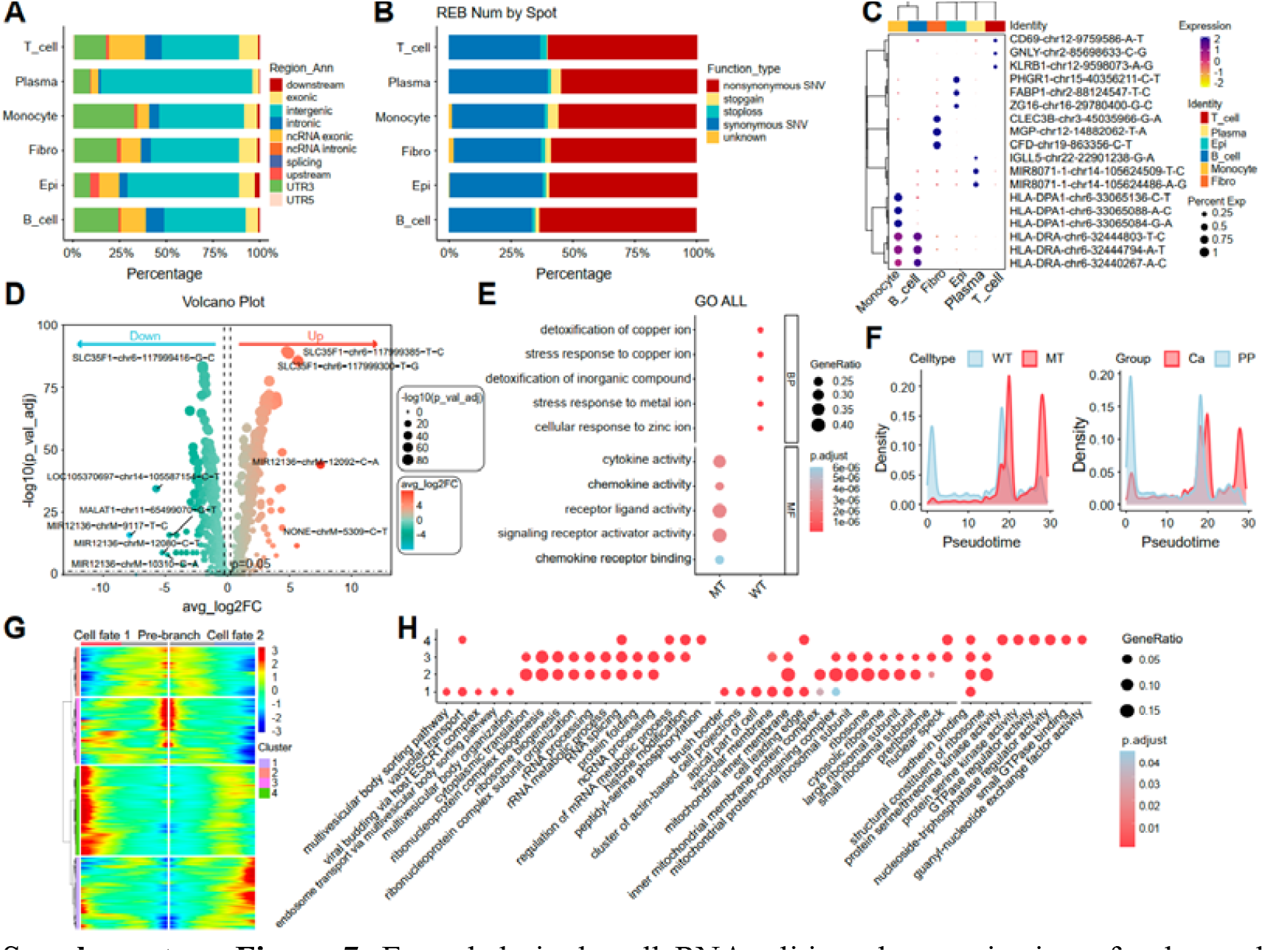
Extended single-cell RNA editing characterization of colorectal cancer. (A) Genomic distribution of RNA editing-like events (3′UTR, exonic, intronic, etc.). (B) Categorization of editing consequences (synonymous, nonsynonymous, etc.). (C) Cell-type-specific RNA editing markers. (D) Volcano plot of differentially edited RNA sites in epithelial cells. (E) GO enrichment of differentially edited genes in epithelial cells. (F) Differentiation states of WT and MT epithelial cells. (G) Gene expression dynamics along the inferred trajectory. (H) Functional pathway enrichment along differentiation states.

**Supplementary Figure 8.**
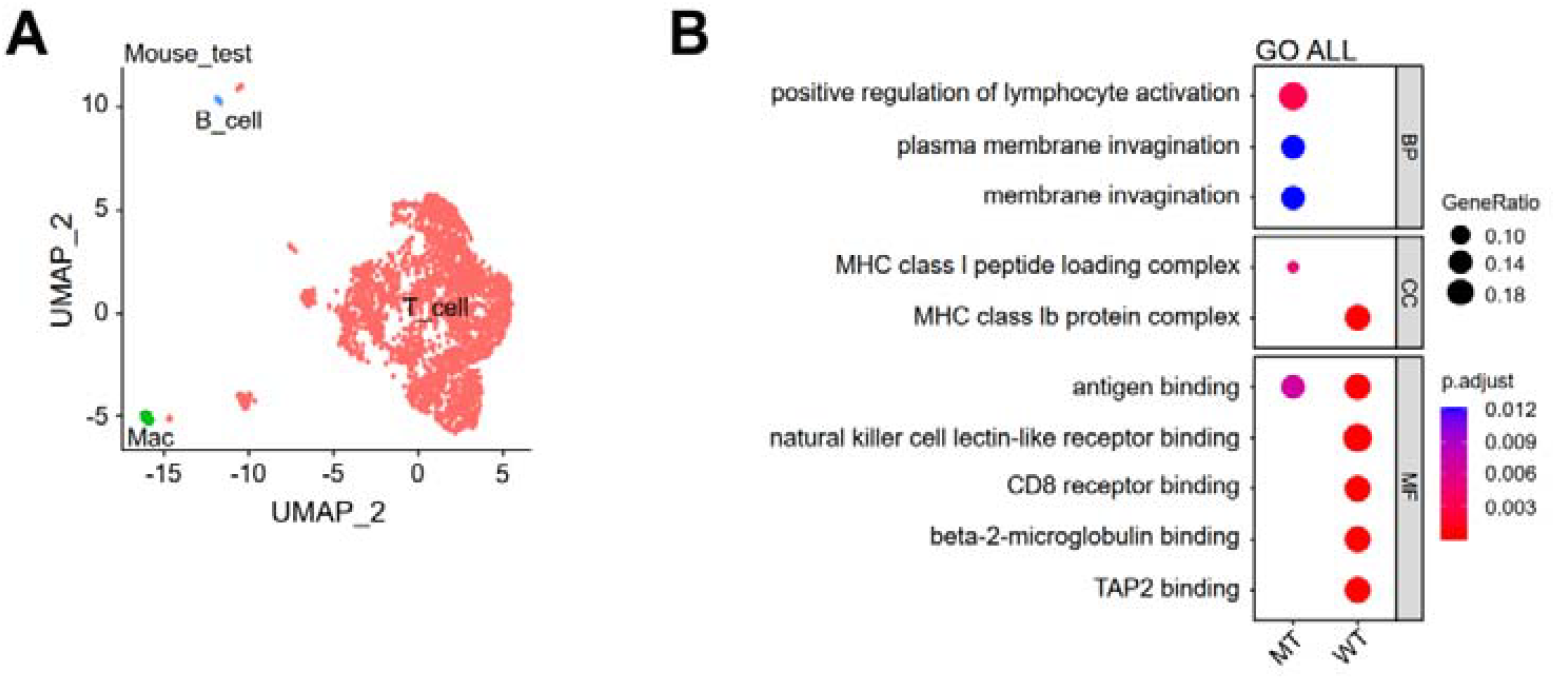
Extended omniRE-seq analysis results of thymic atrophy and recovery in a mouse model. (A) UMAP projection of RNA editing after quality control across all single-cell samples. (B) Gene set enrichment analysis of marker-defined mutant-type (MT) T cell subpopulation.

